# Monitoring in vivo transcription with synthetic serum markers

**DOI:** 10.1101/2024.12.10.627810

**Authors:** Sho Watanabe, Sangsin Lee, Manwal Harb, Shirin Nouraein, Emma Raisley, Honghao Li, Nicolas Buitrago, Beatrice Pforr, Jerzy O. Szablowski

## Abstract

Understanding transcription profiles of living tissues is critical for biology and medicine. However, measurement of the transcript levels is typically done in homogenized tissues post-mortem. Here, we present a new platform that enables non-invasive monitoring of specific mRNA levels *in vivo*, without tissue destruction. We achieved this by combining two cutting-edge tools - synthetic serum markers, called Released Markers of Activity (**RMAs**), and RNA-based sensors of transcription. We call this platform IN-vivo Tracking of ACtive Transcription, or **INTACT**. In INTACT, when the target mRNA is expressed, the RNA sensor detects it and triggers the production and release of RMA reporters into the blood. Once in blood, the RMAs can be easily measured through a simple blood draw. Our data shows that INTACT can measure transcription of transgenes, as well as endogenous transcripts, such as *c-Fos* or *Arc*, both *in vivo* in the brain and in tissue culture. INTACT enables simple measurement of transcript level histories in genetically-targetable cell populations of living animals.

## INTRODUCTION

Transcriptional analysis using techniques such as quantitative polymerase chain reaction (qPCR) or next generation sequencing (NGS) transformed our understanding of biology^1-3^. These techniques usually measure levels of mRNA, an intracellular carrier of biological information which determines the identities^4^, physiological states^5^, and pathogenesis of cells^6-9^. However, available tools measure mRNA levels post-mortem and most cannot record gene expression history or monitor long-term physiological changes in the same animal. The ability to monitor mRNA levels in intact tissues throughout the lifetime of an animal could transform the understanding of *in vivo* systems, similarly to how the qPCR and later NGS transformed such understanding of post-mortem tissue samples.

Monitoring gene expression *in vivo* can be accomplished through noninvasive imaging, such as magnetic resonance imaging (MRI) with genetically-encoded reporters^10-12^, biomolecular ultrasound imaging^13,14,^ or bioluminescence imaging (BLI)^15,16^. However, these techniques use tissue-penetrant forms of energy to measure the reporters, which limits the number of signals that can be monitored at once. Additionally, these methods often contend with signal attenuation in deep tissues, and need expensive bulky equipment that records from immobilized, rather than freely-moving, animals. In particular, the incompatibility of these methods with highly multiplexed readout makes them unlikely to catalyze the kind of leap we observed from monitoring single genes with qPCR, to monitoring thousands of them with transcriptomics. To enable that type of leap, one would first need to convert the levels of mRNA in intact tissues into detectable biochemical signals that are compatible with multiplexed readout.

We recently developed a new type of reporter that enables monitoring of gene expression in deep tissues with a simple blood draw. These reporters, called Released Markers of Activity, or **RMAs**^17^, are proteins that are produced in cells, but released into the blood^18,19^ where they can be easily measured. RMAs can measure transduction and promoter activity in the brain, and by proxy expression of some endogenous genes, such as *c-Fos*. RMAs are highly sensitive, and allowed us to detect their expression in as few as 12 neurons in mice, showed signal levels with up to 100,000-fold over the baseline, and monitored drug-induced neuronal activation^17^. This approach, however, has an important limitation. Relatively few promoters exist that can measure expression of endogenous genes, and fewer still can fit in commonly-used gene delivery vectors^20^. In contrast, in qPCR, one can measure the expression of virtually any gene with an easily-synthesized primer.

Here, we show an approach that is analogous to qPCR, but possible in intact *tissues in vivo*. Similarly to qPCR, this approach can measure levels of theoretically any transcript, but unlike qPCR, it can do so in intact tissues of living animals with a simple blood draw. To achieve this, we combined two cutting-edge technologies – RMAs^17^ (**Figure. 1a, b**), and RNA-based sensors that can detect endogenous mRNA^21-24^ in the cell and in response translate the RMA (**Figure. 1c**). We call this technology, IN-vivo Tracking of ACtive Transcription, or **INTACT**.

**Figure 1.**
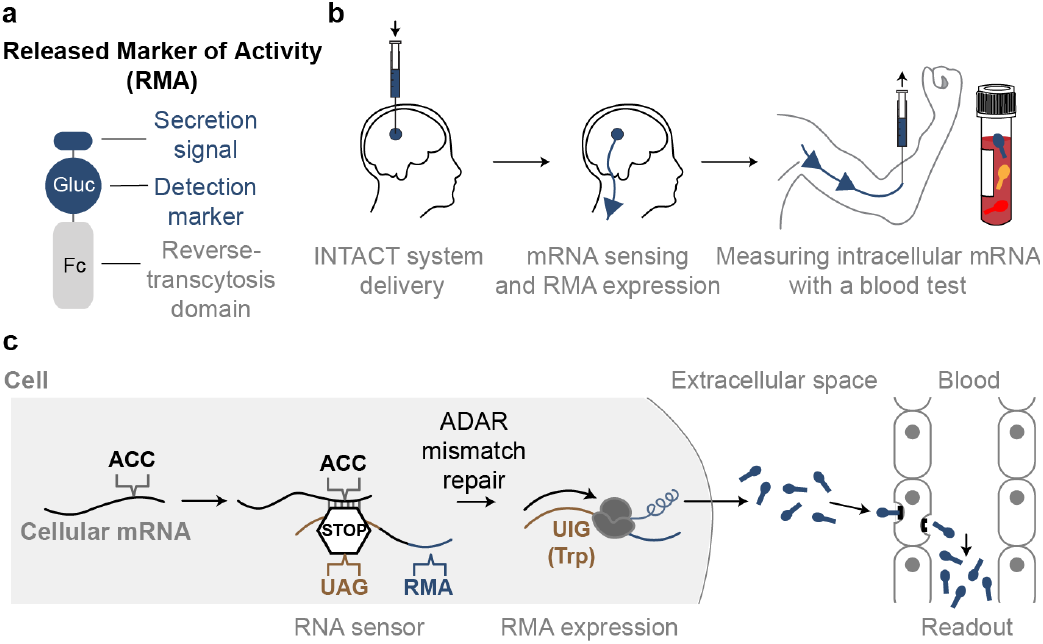
Concept of IN-vivo Tracking of ACtive Transcription (INTACT) **(a)** RMAs are new class of proteins reporters that contain a cell secretion signaling sequence, detectable marker (e.g. luciferase), and a protein fragment that enables reverse transcytosis across the blood brain barrier (BBB), such as the Fc fragment of an antibody. **(b)** INTACT is a new paradigm for monitoring the transcript levels in the brain. Genes encoding INTACT, a synthetic RNA sensor, and a synthetic marker, are delivered to specific brain regions. The INTACT system detects the presence of target mRNA and expresses RMA reporters. RMAs are then secreted from a known brain region into the interstitial space and transported across the BBB into the blood. RMAs can then be detected with biochemical analysis methods allowing with a simple blood test. **(c)** Molecular circuit of INTACT. The RNA sensor is designed to match a specific target mRNA. The sensor includes a stop codon (UAG) and an in-frame reporter gene sequence downstream. When the sensor pairs with the target mRNA, an ADAR enzyme can edit the stop codon (UAG) into a tryptophan codon (Trp). This leads to functional expression of a downstream reporter gene. The amount of reporter produced reflects the level of the target mRNA in the cell.

We decided to prove the concept of INTACT in the brain, due to the sensitivity of the brain to damage and difficulty in its imaging. We have shown that INTACT can be used to track specific neuronal cell-types and monitor neuronal physiology. For mRNA sensing, we utilized a recent paradigm that leverages RNA editing mediated by ADAR (adenosine deaminases acting on RNA)^21-23^ to promote translation of RMA reporters. In these sensors, a short RNA transcript hybridizes with the endogenous target mRNA in a sequence-specific manner, forming a double-stranded RNA (dsRNA), with one mismatched site. That mismatched site contains a stop codon (TAG) within the RNA sensor^25^ that is edited by ADAR into a codon encoding tryptophan, which allows for translation of the sensor to proceed. By placing the RMA sequence downstream of the sensor, we made the expression of RMAs dependent on the presence, or lack thereof, of endogenous mRNA. Consequently, when a sequence-specific intracellular mRNA is produced, RMAs are expressed and released into the blood (**Figure 1c**). The transcription levels of specific mRNA in transduced cell populations are thus linked to blood RMA levels, achieving noninvasive monitoring of mRNA expression in the brain with a simple blood test. We successfully monitored neuronal activation using blood draw by targeting mRNA of immediate early genes (IEG), *c-Fos* and *Activity-regulated cytoskeleton-associated protein* (*Arc*) at different brain regions. We have also shown that INTACT can be used to express RMAs in specific neuronal types, by developing an RNA sensors that are specific to dopaminergic or GABAergic neurons. At last, we demonstrated triplex monitoring of gene expression in one animal by using three orthogonally-detectable RMA reporters. Due to the versatility of the INTACT platform in targeting any gene, its compatibility with the versatile biochemical readout of RMAs, and independence of the readout sensitivity from the tissue depth, INTACT offers a versatile platform to nondestructively study *in vivo* transcript levels.

## RESULTS

### Noninvasive detection of cellular transcripts in tissue culture

To evaluate whether RMAs can report on specific transcript levels, we synthesized an RNA sensor that specifically detects the tdTomato mRNA sequence^21^ and triggers the downstream translation of RMA (RNA sensor^tdT-RMA^). We then transfected plasmids encoding RNA sensor^tdT-RMA^ and tdTomato driven by a CAG promoter (CAG-tdT) into PC12 cells, a widely used murine cell line for studying neurosecretion^26^. Afterwards we measured the amount of secreted RMAs in the culture media using luciferase assays (**Figure. 2a**). We found a significant increase in RMA signals compared to the control group that lacked the target mRNA expression (**Figure. 2b**). The signal fold change between the two groups increased over time (1.35 ± 0.15-fold, P=0.0065 at 12 h, 5.37 ± 0.98-fold, P=0.0003 at 24 h, 26.4 ± 5.89-fold, P=0.0001 at 48 h, 56.4 ± 8.65-fold, P=0.0001 at 72 h, and 83.0 ± 16.6-fold, P=0.0002 at 96 h over the baseline, using two-tailed unpaired t-test using *Benjamini– Hochberg* for false discovery rate (FDR) adjustment). The levels of secreted RMA signal correlated with the expression levels of tdTomato mRNA, as determined by qPCR, following a Michelis-Menten curve – presumably due to the translation of RMA being catalyzed enzymatically by ADAR (R=0.987, **Figure. 2c**). Consistent with previous studies^21-23^, we observed that overexpression of exogenous ADAR (ADAR2 or p150 isoform of ADAR1) increased the mean RMA levels by 183 ± 63% for ADAR2 (P=0.0002) and 175 ± 98% for ADAR1p150 (P=0.0002), respectively (one-way ANOVA with Dunnett’s test, **Extended Data Fig. 1**).

**Figure 2.**
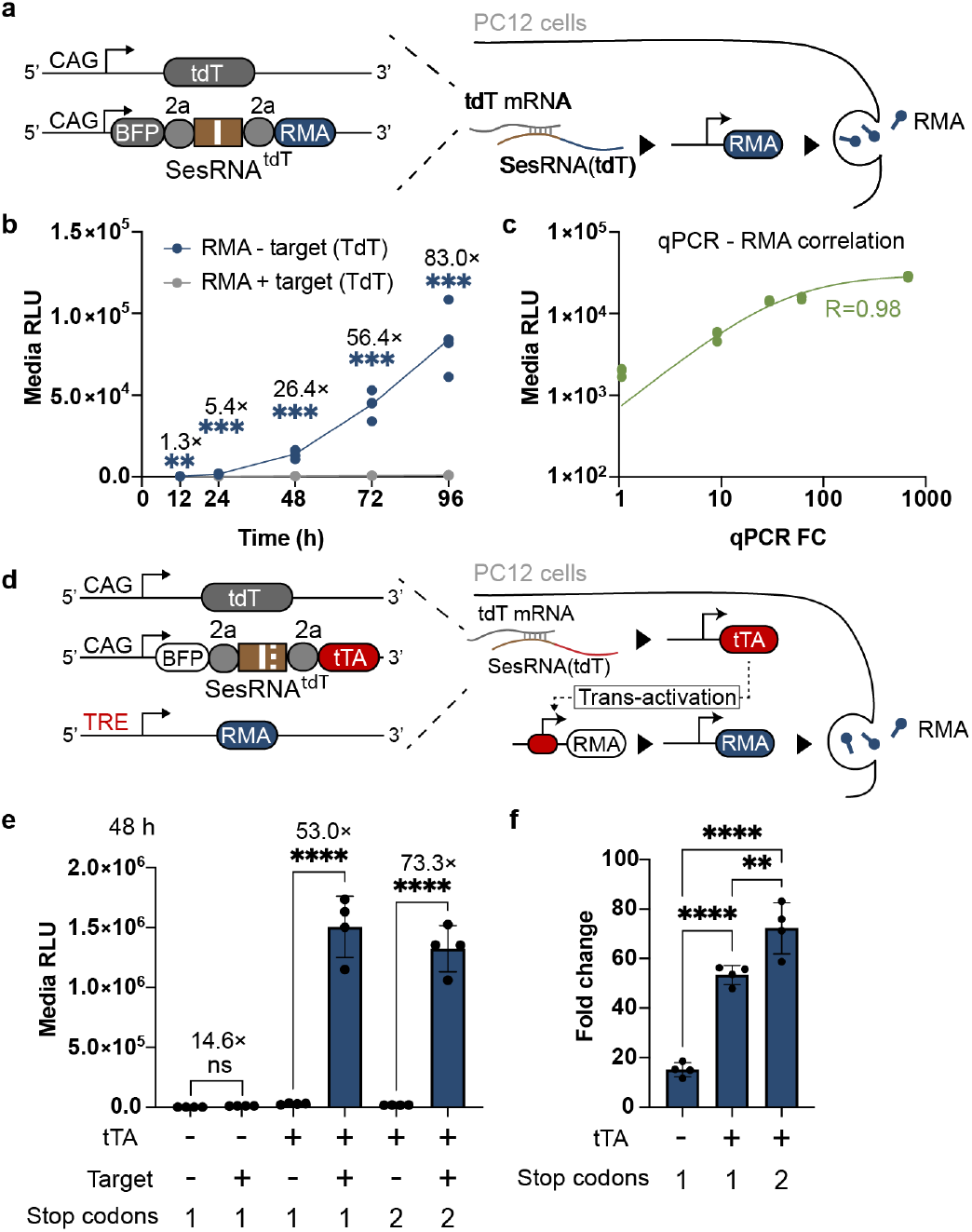
Intracellular transcript monitoring with RMA in tissue culture. **(a)** Design of the RNA sensor circuit. CAG-tdT encodes a target mRNA (tdTomato). RNA sensor encodes a BFP sequence followed by tdTomato-targeting sequence (SesRNA^tdT^) and the RMA. SesRNA^tdT^ has a reverse complementary sequence that can hybridize with the tdTomato mRNA. The white line in SesRNA^tdT^ indicates the presence of a stop codon. The ‘2a’ indicate RNA sequences encoding a self-cleaving peptide. **(b)** Levels of RMA in the cell culture medium in response to tdTomato transduction. N=4 independent cultures were analyzed. In comparison of each group with and without the transfection of CAG-tdT, the signal fold change between the two groups increased over time (1.35 ± 0.15-fold, P=0.0065 at 12 h, 5.37 ± 0.98-fold, P=0.0003 at 24 h, 26.4 ± 5.89-fold, P=0.0001 at 48 h, 56.4 ± 8.65-fold, P=0.0001 at 72 h, and 83.0 ± 16.6-fold, P=0.0002 at 96 h over the baseline, using unpaired two-tailed multiple t-test with the *Benjamini-Krieger-Yekutieli* false discovery rate (FDR) adjustment (5%). **(c)** Mean RMA signals (n=3, biological replicates) are plotted against the tdTomato gene expression levels as determined by qPCR. Target tdTomato expression range is created by using different amount of plasmid DNA (1, 10, 50, 100, 500 ng for 100,000 cells) for transfection. The value of qPCR fold change (FC) at each group of expression level is determined as a mean of n=3 biological replicates. The green line represents the linear regression results of the data. *R*-value denotes correlation between qPCR and medium RMA signals fitted to Michelis-Menten kinetics curve. **(d)** Design of the tetracycline-operated sensor circuit. The tetracycline transactivator (tTA2) is expressed when the stop codon is removed from tdTomato binding region of the RNA sensor. The tTA2 then activates the TRE promoter to drive RMA expression. The white lines in SesRNA^tdT^ indicate the presence of one or two stop codons depending on the construct. **(e)** The tTA2-TRE-based sensor shows significant increase in production of RMAs in the presence of the tdT (target) mRNA expression. n=4 independent cultures were analyzed. n.s. in comparison of the group of non-tTA sensor (Fig. 2a) and P<0.0001 in comparison of other groups with and without the transfection of CAG-tdT, using two-way ANOVA (F_3.9_=138.2) with Tukey’s test. **(f)** Influence of the sensor design on the RMA expression levels. The sensor using tTA2-TRE-based amplification and two stop codons leads to the highest fold change between cells expressing tdT mRNA and vehicle controls. n=4 independent cultures were analyzed. P<0.0001 for the comparison of a single stop codon with and without the tTA2-TRE circuit; P=0.007 for the comparison two tTA2-TRE sensors with a single or two stop codons; P<0.0001 for comparison between the sensor with a single stop codon and no tTA2 amplification, and the sensor with two stop codons and tTA2-TRE amplficiation. P-values obtained using one-way ANOVA (F_2,9_=79.33, P<0.0001) with Tukey’s test. ^**^P<0.01, ^***^P<0.001, and ****P<0.0001. All data are shown as mean ± s.d.

To enhance detection sensitivity, we engineered a tetracycline-operated sensor circuit containing an RMA reporter gene driven by the TRE3g promoter (TRE3g-RMA) and a tdTomato-targeting RNA sensor expressing tTA2 (RNA sensor^tdT-tTA2^) (**Figure. 2e**). tTA2 is produced in the presence of tdTomato mRNA and drives expression of RMA from a TRE3g promoter^21,27^. We co-transfected RNA sensor^tdT-tTA2^, TRE3g-RMA, and CAG-tdT and saw 53.0 ± 7.80-fold change (P<0.0001) over the baseline at 48 h compared to the control group lacking CAG-tdT. On the other hand, Sensor without tTA2-based amplification showed 14.6 ± 0.87-fold change (N.S., P>0.05) over the baseline at 48 h, compared to the group lacking CAG-tdT. (two-way ANOVA with Tukey test, **Figure. 2e**). Furthermore, previous research showed that increasing the number of stop codons at the complementary binding sequence of RNA sensors reduces sensor translation in the absence of the target^21^. We therefore incorporated two stop codons in our RNA sensor and observed a reduction of the baseline RMA expression, resulting in an enhanced signal change of 72.3 ± 9.14-fold over the baseline at 48 h (P<0.0001, two-way ANOVA with Tukey test, **Figure. 2e**). The differences in fold changes over the baseline were significant between the three types of RNA sensors (one-way ANOVA with Tukey test, **Figure. 2f**).

To examine whether RMAs are successfully released from cells or accumulate within them, we observed the intracellular localization of RMA immunofluorescence signal under confocal microscope. HEK293T cells expressing CAG promoter-driven RMA (CAG-RMA) showed fluorescence patterns sequestrated primarily in the periphery of the cell (**Extended Data Fig. 2a**). However, cells expressing RNA sensor^tdT-RMA^ and CAG-tdT showed more pronounced nucleic and cytosolic localization patterns (**Extended Data Fig. 2a**). We hypothesized that when the ribosome encounters the auto-cleavage P2A/T2A sequence in RNA sensors, the following mRNA encoding the RMA reporter fails to be recruited to endoplasmic reticulum ER^28^, leading to partial translation of RMAs in the cytosol, and interference with membrane localization and secretion. To avoid this potential effect and to maximize the RMAs’ release, we decided to use sensor^tdT-tTA2^ for subsequent *in vivo* experiments. The sensor^tdT-tTA2^ translates tTA2 which, can then drive expression of RMA without any potential effects of P2A/T2A in cis configuration. Supporting this hypothesis, we found that cells expressing the RNA sensor^tdT-tTA2^, TRE3g-RMA, and CAG-tdT showed more prominent localization near the cell periphery than when expressing the sensor^tdT-RMA^ alone (**Extended Data Fig. 2a**). To quantify these microscopy results, a blinded observer scored the cells for presence of a primarily peripheral RMA immunostaining (score of 2), comparable cytoplasmic and peripheral signal (score of 1), or undetectable peripheral signal (score of 0). We found significant differences between each group (P<0.0001 for all groups, one-way ANOVA with Tukey test, **Extended Data Fig. 2b**).

### Detection of exogenous gene expression using INTACT in mice

To test whether INTACT could monitor mRNA expression *in vivo*, we constructed an RNA sensor that targets *tdTomato* mRNA and expresses tTA2 upon activation to drive RMA expression from TRE3G promoter (**Figure 3a**). The sensor was expressed under the human synapsin (hSYN) promoter to restrict the monitoring to neurons. Further, to account for variations in gene delivery efficiency between mice, each mouse included an internal control - an independently-detectable *Cypridina luciferase* (Cluc)-RMA, which was constitutively expressed under the hSyn promoter^17^. Overall, each mouse received AAVs encoding: 1) hSyn1-EGFP-Sensor RNA^tdT-tTA2^, 2) TRE3g-RMA, 3) CAG-tdT, and 4) hSyn1-Cluc-RMA through a direct intracerebral injection into the left caudate putamen (CP). Two and three weeks after transduction we collected the blood and measured the plasma RMA signals (**Figure. 3a**). We found that when *tdTomato* mRNA was expressed under the CAG promoter, the plasma RMA signals were significantly higher than the baseline by 18.8 ± 1.87-fold (P<0.0001) at two weeks and 10.4 ± 1.99-fold (P<0.0001) at three weeks post-transduction (two-tailed unpaired t-test, **Figure. 3b**). In post-mortem histology we observed 89.0 ± 2.61% specificity of tdTomato transcript detection, as shown by the fraction of cells with positive RMA staining and concurrent tdTomato fluorescence (**Figure 3c, d**). We observed positive RMA staining among 61.4 ± 5.12% of the cells that expressed both the target (tdT) and the sensor (GFP) (**Extended Data Fig. 3**), which is comparable to the previously reported efficiency of sensing for the tdTomato sensor driving eGFP in tissue culture^21^. We also observed some neurons expressed RMA proteins without the tdTomato expression, possibly due to the unexpected stop codon readthrough in the RNA sensor^29^ or nonspecific leakage of the TRE promoter^30^.

**Figure 3.**
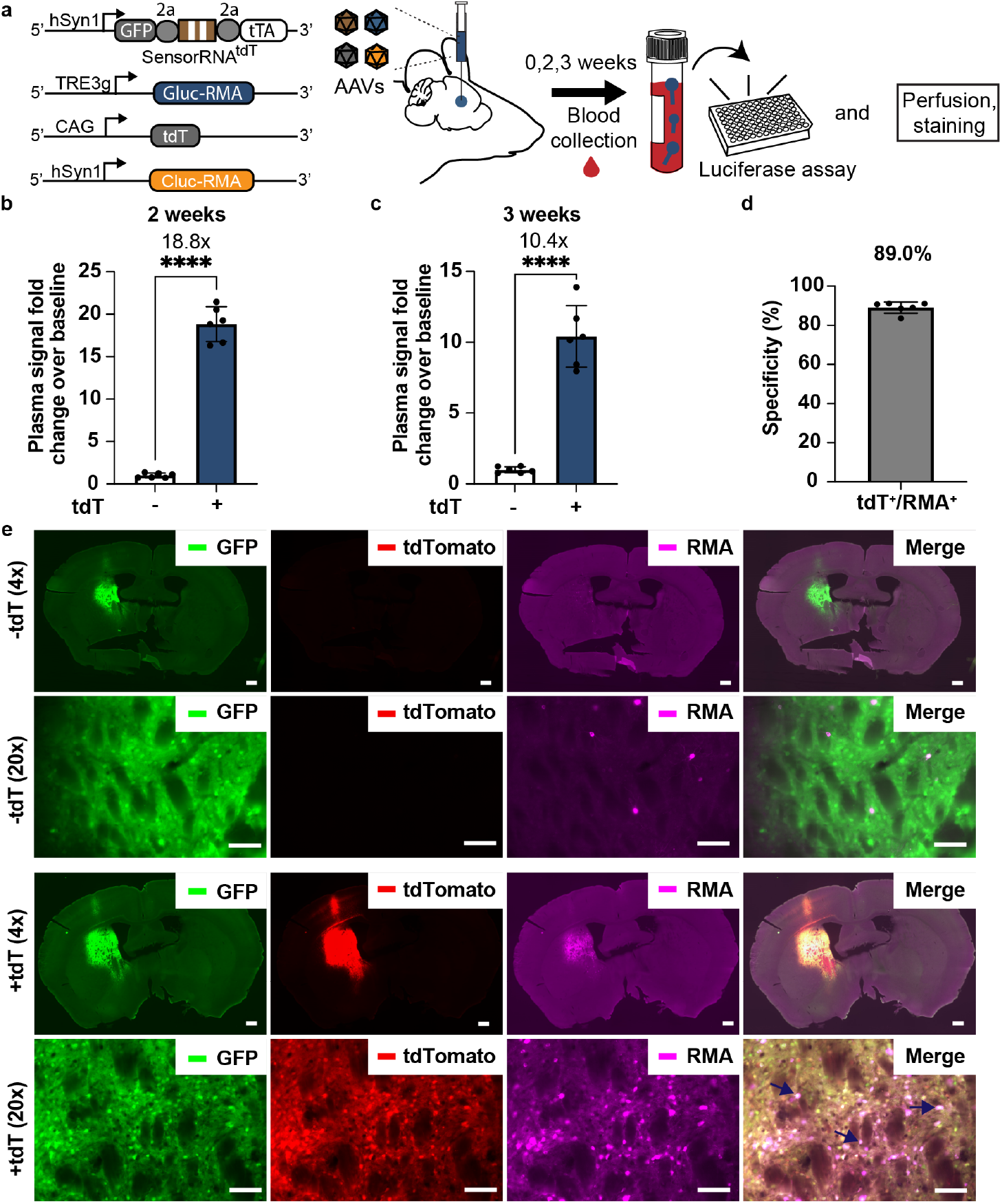
Detection of transgene mRNA with INTACT *in vivo*. **(a)** Experimental design for testing INTACT in mice. Collection of the baseline blood sample was followed by injection of four different AAVs encoding the RNA sensor of tdT, TRE-Gluc-RMA reporter, CAG-tdT, and hSyn1-Cluc-RMA into the brain parenchyma (0.9×10^9^ vg for each AAV). Afterwards, blood collection was performed at 2 and 3 weeks. **(b)** Plasma bioluminescence signal to evaluate RMA expression at two weeks, and **(c)** three weeks after AAV injection. Control group lacked the AAV encoding CAG-tdT. Gluc-RMA signal is normalized to the Cluc-RMA to compensate for the variation in transduction between mice. p<0.0001 for comparison of each group (n=6) with and without the transfection of CAG-tdT, using unpaired two-tailed t-test. **(d)** The specificity of tdT-target RNA sensor was assessed by counting the fraction of RMA-positive cells that also expressed tdT. n=6 independent mice were analyzed. **(e)** Representative images showing gene expression of RNA sensor (GFP), target mRNA (tdTomato), and RMAs. The arrows indicate the cells that express RNA sensor, tdTomato, and RMA-positive cells. ^****^P<0.0001. All data are shown as mean ± s.d. Scale bar, 500 μm for whole brain (4x) images and 100 μm for magnification (20x) images.

### Noninvasive monitoring of gene delivery to specific cell types with INTACT

The brain contains hundreds of neuronal subtypes^31^, but only some are accessible with promoters that can fit within AAV packaging capacity^32^. We decided to apply INTACT to restrict expression of RMAs to specific neuronal subtypes, starting with targeting of dopaminergic neurons, which can only be targeted *in vivo* with a long 2.5kb *tyrosine hydroxylase* (*TH*) promoter that covers more than 50% of the AAV packaging capacity^33^. To identify the best TH sensor sequence, we tested several variants in tissue culture and identified the one that generated the highest reporter signals in PC12 cells (**Supplementary Fig. 1**). We packaged this sensor in AAV and expressed it constitutively under hSYN promoter. The sensor produced tTA2 upon TH-mRNA detection to drive the expression of Gluc-RMA. As in other experiments, as an internal control, we also co-administered AAV.php.eb carrying Cluc-RMA expressed under hSYN promoter. All constructs (**Figure. 4a**) were co-injected into the ventral tegmental area (**VTA**), a region which contains TH-positive neurons. At two and three weeks after AAV delivery, plasma Gluc-RMA signals were 587 ± 98.6-fold (P<0.0001) and 390 ± 150-fold (P=0.0005) higher than the baseline at 0 weeks, respectively, as shown by one-way ANOVA with Tukey test (**Figure 4b**). Histological analysis revealed that Gluc-RMA expression was specific to TH mRNA-positive cells at the local injection site (**Figure 4c**), with 93.9 ± 3.36% of RMA-positive cells also being TH positive (**Figure 4d**).

**Figure 4.**
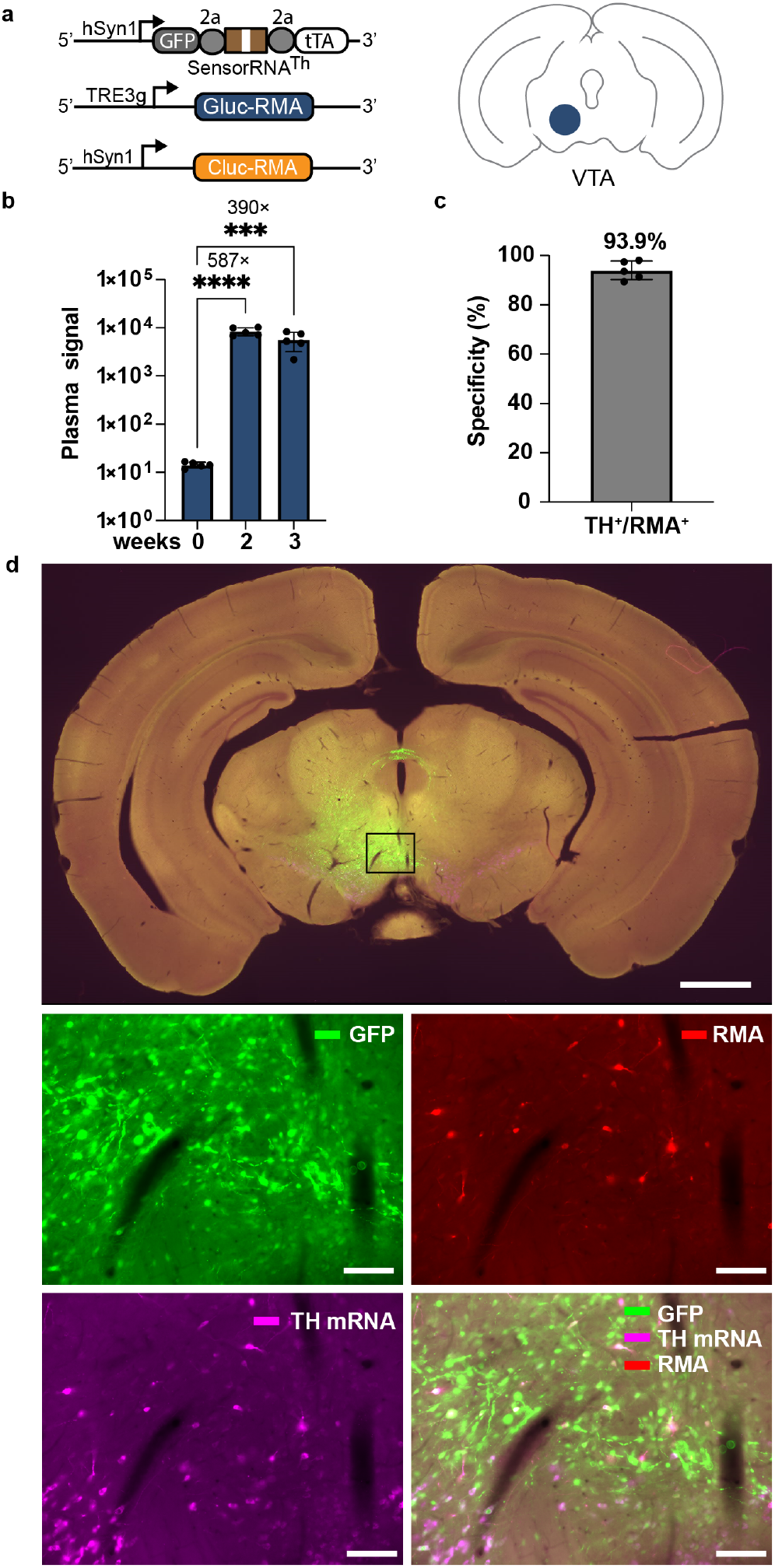
Monitoring transduction of specific neuronal subtypes in the brain. **(a)** Experimental design for detection of endogenous mRNA with INTACT. Baseline blood sample was collected, and three different AAVs (0.9×10^9^ vg for each AAV) encoding Th-targeting RNA sensor, TRE-Gluc-RMA reporter, and hSyn1-Cluc-RMA were injected into the ventral tegmental area (VTA). Then, the blood was collected at 2 and 3 weeks after the AAV delivery. **(b)** Plasma bioluminescence signals were higher at two (P<0.0001) and three weeks (P=0.0005) time points compared to the 0-week baseline. n=5 independent mice were analyzed using one-way ANOVA (F_2,12_=33.00, P<0.0001) with Tukey’s test. **(c)** The specificity of Th-targeting RNA sensor was assessed by counting the fraction of RMA-positive cells that also expressed TH. n=5 independent mouse data were analyzed. **(d)** Representative images showing gene expression of RMA sensor (green), *TH* mRNA (magenta), and Gluc-RMA (red). Gluc-RMA was labeled by Gluc antibody. Endogenous *TH* mRNA was labeled by in situ hybridization using *TH* probes. ^***^p<0.001, ^****^p<0.0001, Scale bars 1000 μm and 100 μm of the top (whole brain) and below (enlarged) images, respectively. Data is shown as mean c s.d.

To monitor different neuronal subtypes, we synthesized the hSyn1-controlled *Vesicular GABA Transporter* (*Vgat*) mRNA-targeting RNA sensor by using a previously published *Vgat* mRNA-targeting sensor^21^ for expression of RMAs. We delivered the sensor into the mouse striatum along with the TRE3g-RMA reporter and hSyn1-Cluc-RMA (**Extended Data Fig. 5a**). At 2 weeks and 3 weeks after AAV delivery, plasma Gluc-RMA signals were 1295 ± 507.2-fold (P=0.0004) and 907 ± 288-fold (P=0.0065) higher than the baseline at 0 weeks, respectively (one-way ANOVA with Tukey test, **Extended Data Fig. 5b**). Similar to the *TH* detection (**Figure. 4**), the expression of Gluc-RMA was specific to *Vgat* mRNA-positive cells in 94.1 ± 2.38% cells (**Extended Data Fig. 5c, d**). Together, these data provide evidence that Gluc-RMA reporters can monitor expression of cell-type marker mRNA in specific neuronal populations with a blood test.

### INTACT can monitor transcription of immediate early gene (IEG) c-Fos in vivo

To demonstrate the ability of INTACT platform to monitor sustained neuronal activity, we next sought to record mRNA levels of the immediate early genes (IEGs). We chose *c-Fos* and *Activity Regulated Cytoskeleton Associated* (*Arc)* proteins, both of which are expressed during heightened neuronal activity^34^. We induced the neuronal activity using an excitatory designer receptor exclusively activated by a designer drug (DREADD) - hM3Dq^35^ that was co-injected into the brain along with INTACT components. The hM3Dq elicits neuronal firing and c-Fos accumulation upon drug-induced activation^35,36^.

We first designed the *c-Fos* mRNA and *Arc* mRNA targeting sensors and screened four or five variants of these sensors in tissue culture in HEK293 cells by co-expressing the candidate sensors with their target *c-Fos* or *Arc* mRNA sequences (**Supplementary Fig. 2, 3**). We then placed these sensors before a downstream tTA2, and under the control of hSyn promoter to restrict IEG sensing to neurons (**Figure 5a**). We then co-injected AAVs carrying the sensor, RMA under TRE3g promoter, hSyn1-Cluc, and DREADD AAVs into the striatum of the brain for *c-Fos* mRNA monitoring (**Figure 5a**). To record a discrete time window, mice were placed on a doxycycline (Dox) chow diet throughout the AAV incubation period to prevent tTA2 from binding to the TRE promoter and expressing RMA. Mice went on a Dox-free diet 48 h before administering CNO to enable recording of neuronal activation (**Figure 5a**).

**Figure 5.**
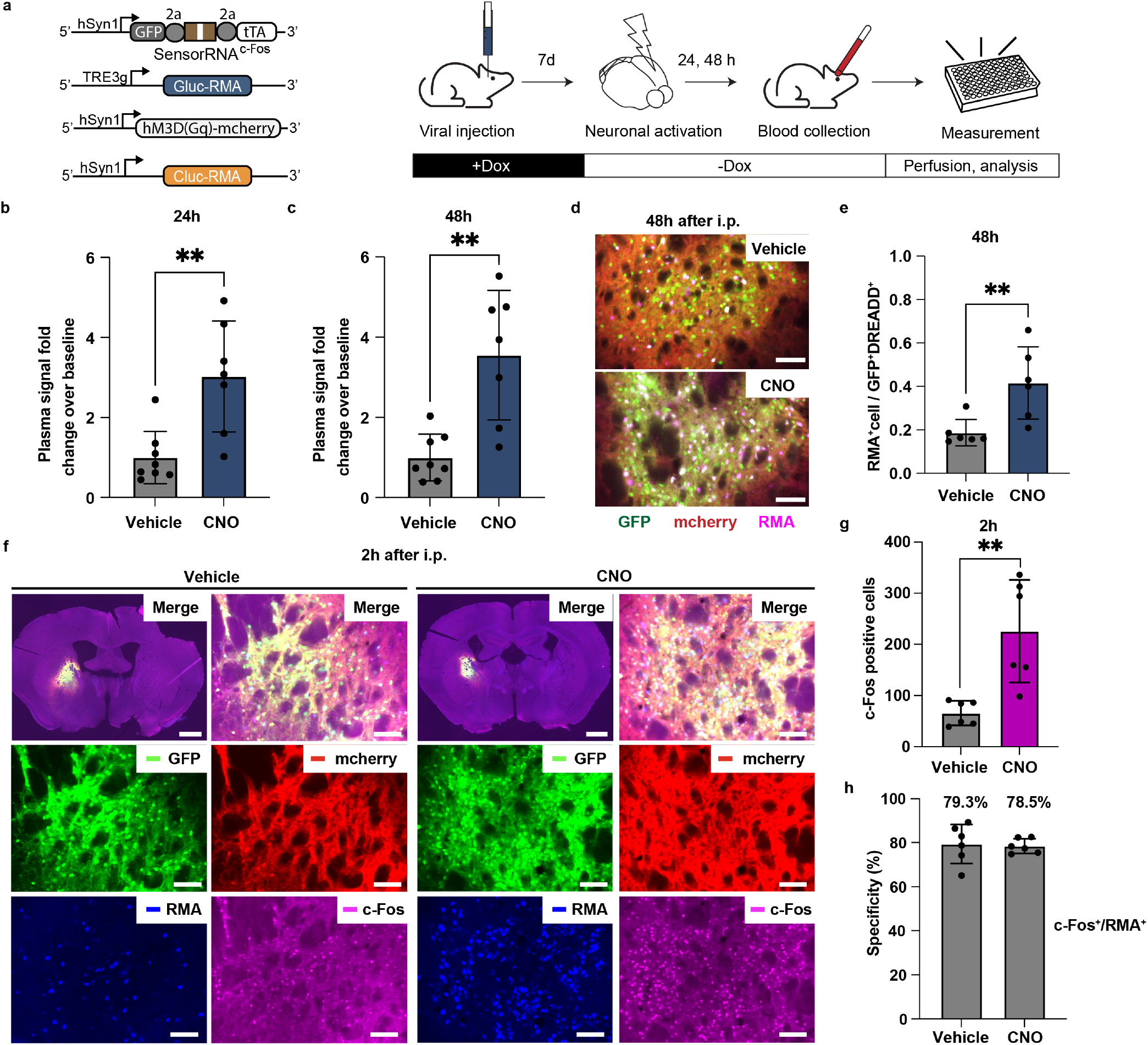
INTACT detects neuronal activity in vivo. **(a)** Monitoring chemogenetic activation of neurons with INTACT. RNA sensor was designed to target endogenous *c-Fos* mRNA, a marker of elevated neuronal activity. Mice underwent injection of AAVs encoding RNA sensor (Syn-sesRNA(c-Fos)-tTA2), TRE-Gluc-RMA, activatory DREADD (hM3Dq) receptor, and hSyn1-Cluc-RMA. Mice were then placed on a doxycycline (Dox) diet which prevents tTA2 from binding to TRE promoter, and thus prevents the expression. This design allows for temporal gating the recording of neuronal activity. The Dox diet was then withdrawn, and 48 hours later one group received CNO to activate DREADD-expressing neurons, while the other group received a vehicle control. Blood samples were collected afterwards to measure Gluc-RMA levels. **(b)** Gluc-RMA signal divided by the pre-CNO injection baseline at 24 h, and **(c)** 48 h after CNO administration. Signals were normalized by dividing the RLU values obtained from Gluc-RMA over Cluc-RMA. n=8 (vehicle, 0 mg kg^-1^) and n=7 (+CNO group, 2.5 mg kg^-1^) independent mice were analyzed using unpaired two-tailed t-test at 24 h (P=0.0022 (b) and 48 h (P=0.0011) (c) time points. ^**^p<0.01. **(d)** Representative images of gene expression of RNA sensor (GFP), hM3Dq (mCherry), and Gluc-RMA (Magenta) in the brain section at 48 h after CNO (2.5 mg kg^-1^) or vehicle injection. Scale bar, 500 μm. **(e)** Fraction of cells expressing RMAs among those that also showed transduction of the sensor (GFP) and DREADD (mCherry) in post-mortem histology, 48 hours after CNO or vehicle injection (n=6 mice in each group). P=0.01 using unpaired two-tailed t-test. ^**^P<0.01. **(f)** Representative images showing gene expression of RNA sensor (GFP), DREADD Gq (mCherry), Gluc-RMA (blue), and c-Fos (magenta) at 2 h after i.p. Scale bars 1000 μm and 100 μm of the top left (whole brain) and the others (enlarged) images, respectively. **(g)** The number of c-Fos expressing cells in the targeted region at 2 h after injection of CNO or vehicle (n=6 mice in each group). P=0.0030 using unpaired two-tailed Student’s t-test. ^**^P<0.01. **(h)** The specificity of c-Fos targeting RNA sensor. Specificity was assessed by counting the number of cells expressing all components of INTACT that also were positive for c-Fos (n=6 mice in each group). Data is shown as mean ± s.d.

We found that mice injected with CNO (2.5 mg kg^-1^) had plasma Gluc-RMA signals that were 3.03 ± 1.28-fold and 3.55 ± 1.49-fold higher than those injected with vehicle at 24 and 48 h after activation, respectively (P=0.0026, and P=0.0011, respectively, using two-tailed unpaired t-test, **Figure 5b, c**). After 48 hours we perfused the animals to analyze the brain tissue histologically. We chose top 6 mice with highest level of transduction from each group for histological analysis, as determined by serum Cluc control levels. We aimed to measure the responsivness of RMA to c-Fos, while ensuring quantification was performed in successfully and equally transduced mice. Cluc-levels were identical for each group before CNO/vehicle administration, indicating comparable transduction (P=0.494, two-tailed unpaired t-test). We found that RMAs were produced preferentially in c-Fos positive cells. The number of cells expressing RMAs in cells containing both RNA sensors (GFP) and DREADD receptor (mCherry) was 2.45 ± 0.53-fold higher in the CNO group than in the vehicle group (P=0.01, two-tailed unpaired t-test, **Figure 5d, e, Supplementary Fig. 4**). To confirm the DREADDs were inducing c-Fos in our samples, we analyzed the brain tissues two hours after neuronal activation, as previously published^35^. As expected, CNO group showed higher c-Fos positive cell numbers compared to the vehicle group suggesting successful chemogenetic activation (3.44 ± 1.40-fold, P=0.0030, two-tailed unpaired t-test, **Figure 5f, g**). The specificity analysis showed 79.4 ± 8.08% and 78.5 ± 3.03% of RMA-expressing neurons also expressed c-Fos protein within both the vehicle and CNO-injected groups, respectively (**Figure 5f, h**).

Following the demonstration of *c-Fos* mRNA monitoring we next injected the AAVs constituting INTACT into the striatum and hippocampus CA1, respectively, to monitor *Arc* mRNA (**Extended Data Fig. 6a**). Exploration of new environments was shown to induce Arc expression in the hippocampus^37^. On the other hand, the striatum showed regions-specific patterns of *Arc* expression under the training of inhibitory avoidance^38^, suggesting both of these regions are capable of *Arc* induction. Indeed, we found Arc expression in both of these regions, however, it was more strongly activated in CA1 of the hippocampus compared to CP of the striatum after DREADD-induced neuronal activation (P=0.0001, two-tailed unpaired t-test, **Extended Data Fig. 7a–d**).

Our results showed that mice injected with CNO (2.5 mg kg^-1^) generated plasma Gluc-RMA signals that were 1.26 ± 0.21-fold and 4.25 ± 1.51-fold higher than those injected with vehicle at 24 and 48 h after activation, respectively (P=0.0317, and P=0.0026, respectively, two-tailed unpaired t-test, **Extended Data Fig. 6b, c**). After 48 hours, we perfused the animals to analyze the brain tissue histologically. We found more RMA-expressing cells among those cells that also contained both RNA sensors (GFP) and DREADD receptor (mCherry) in the CNO group than in the vehicle group (2.45 ± 0.53-fold, two-tailed unpaired t-test, p<0.0001, **Extended Data Fig. 6d, e**). To confirm *Arc* induction by DREADD, we analyzed the brain tissues two hours after neuronal activation. The group administered with CNO exhibited higher Arc-positive cells compared to the vehicle group, suggesting the observed increase in plasma RMA signals (19.3 ± 4.73-fold, P=0.001, two-tailed unpaired t-test, **Extended Data Fig. 6f, g**). We could not calculate the specificity of *Arc* sensor since there were few RMA-positive cells observed at this early time point (**Supplementary Fig. 5**). We reasoned that RMA protein might not have been translated within the short 2 h timeframe in response to *Arc* mRNA sensing, as was previously the case with the c-Fos promoter^17^.

Contrary to the data in the hippocampus, we could not observe significant increases of plasma Gluc-RMA signals in mice injected with CNO (2.5 mg kg^-1^) in which AAVs were injected into the striatum, both at 24 and 48 h after neuronal activation (N.S. P>0.05, two-tailed unpaired t-test, **Extended Data Fig. 6h, i**). In histology, the number of RMA-positive cells was also not significantly different between the CNO group and the control group (P=0.7038, two-tailed unpaired t-test, **Extended Data Fig. 6j, Supplementary Fig. 6**), which was consistent with the data of serum RMA signal. These data suggested that Gluc-RMA might not be sensitive enough to capture the lower levels of Arc expression in the striatum (**Extended Data Fig. 7a–d**). Thus, we sought to develop a more sensitive RMA reporter to enable monitoring of *Arc* in the striatum.

### Multiplex monitoring of gene expressions by using orthogonal RMA reporters

We aimed to extend the versatility of INTACT system toward multiplex detection. One of the advantages of INTACT is the detection of marker by any chosen biochemical analysis, which has the potential for highly multiplexed of readout. As a starting point, we sought to demonstrate detection of three different markers based on luciferases: Gluc, Cluc, and Nanoluc (Nluc)^39^. We synthesized them (**Extended Data Fig. 8a**) and confirmed their secretion and orthogonal detection in vitro with different substrates that are specific to each luciferase (one-way ANOVA with Tukey test, **Extended Data Fig. 8b–f**). Nluc-RMA showed 2.1 ± 0.6-fold higher signal fold changes over the baseline compared to Gluc-RMA (P=0.013, two-tailed unpaired t-test) when targeting the same target mRNA at the last commonly tested timepoint, 72 h (**Figure 2b, Extended Data Fig. 8g**). Due to the higher sensitivity of Nluc-RMA, we hypothesized that Nluc-RMA had a potential to detect *Arc* expression in the striatum in response to neuronal activation. We further hypothesized that the three luciferases could be detected within the same animal to monitoring neuronal activation simultaneously.

To test these hypotheses, we injected AAVs encoding *Arc* sensor under hSyn promoter with a downstream tTA2, along with the TRE3g-Gluc-RMA reporter into the hippocampus CA1 and the same *Arc* sensor, along with the TRE3g-Nluc-RMA reporter into the striatum (**Figure 6a**). At the same time, we injected AAV encoding hSyn1-Cluc-RMA into the substantia nigra in the midbrain as a negative control unaffected by DREADD induction. As previously, we set up a discrete monitoring time window, controlled by Dox chow diet and withdrew Dox 48 h before administering CNO (**Figure 6a**) to begin monitoring *Arc* expression.

**Figure 6.**
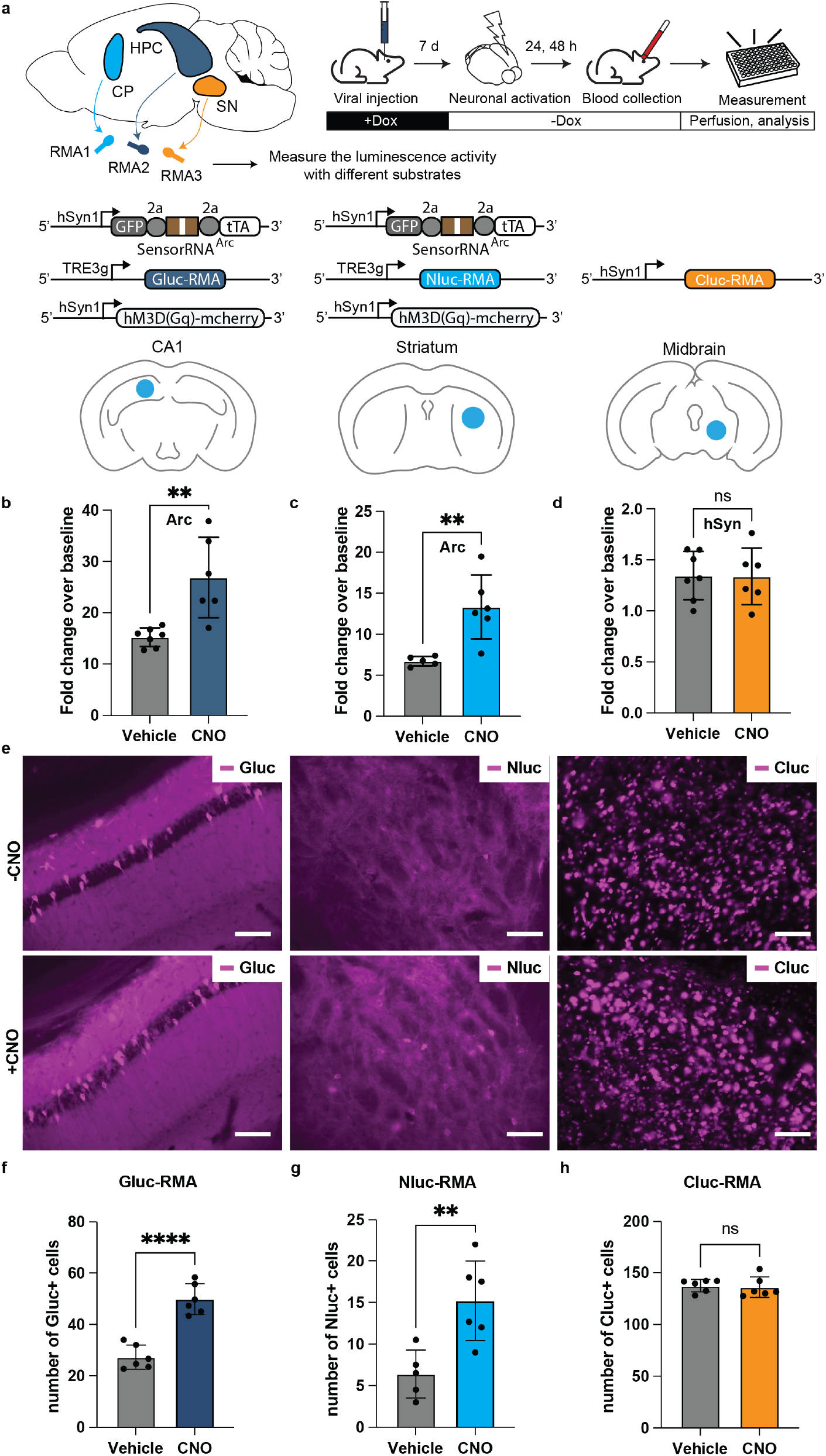
Multiplex monitoring of Arc expression with orthogonal RMA reporters. **(a)** Monitoring gene expression in multiple brain regions with INTACT. AAVs encoding RNA sensor for *Arc* mRNA were injected into the hippocampus CA1 and striatum of mice. Each of these injections also contained AAVs encoding TRE-Gluc-RMA (CA1) or TRE-Nluc-RMA (striatum) and the activatory DREADD receptor under hSYN1 promoter (hM3Dq). Additionally, an AAV encoding hSYN1-Cluc-RMA was injected into the substantia nigra of the midbrain. Mice were then placed on the Dox diet to shut down the RMA production until the desired timepoint for *Arc* recording. After 7 days, the Dox diet was withdrawn, and 48 hours later mice received either CNO or vehicle injections. Blood was then drawn at 24h and 48h to analyze the serum RMA levels. **(b)** Relative fold change of plasma bioluminescence over 0 h baseline of Gluc-RMA, **(c)** Nluc-RMA, and **(d)** Cluc-RMA at 48 h post-CNO or vehicle administration (n=7 mice in vehicle and n=6 mice analyzed in the CNO groups; P=0.0028 for Gluc, P=0.0048 for Nluc, and P=0.9532 for Cluc RMAs using two-tailed unpaired t-test). **(e)** Representative images showing RMA expression (magenta) in the brain at 48 h after vehicle or CNO administration. The expression of Gluc-RMA and Nluc-RMA was confirmed by immunostaining, while Cluc-RMA was detected by fluorescent in situ hybridization. Scale bar, 500 mm. **(f-h)** The number of RMA-positive cells in the targeted brain region 48 h after CNO or vehicle administration. For Gluc- and Cluc-RMAs n=7 (vehicle) and n=6 (CNO) mice were analyzed, while Nluc-RMA used n=5 (vehicle) and n=6 (CNO) mice were analyzed using unpaired two-tailed t-test (P<0.0001 for Gluc-RMA **(f)**, P=0.0059 for Nluc-RMA **(g)**, P>0.05 for Cluc-RMA **(h)**). ^****^P<0.0001, ^**^P<0.01, ns P>0.05. Data is shown as mean ± s.d.

We collected blood before, 24 h and 48 h after CNO (2.5 mg kg^-1^) injection, and measured the luminescence activity by adding the three different substrates, respectively. We observed that mice injected with CNO generated plasma Gluc-RMA signals, which are derived from *Arc* sensing at hippocampus CA1, that was 1.76 ± 0.47-fold (P=0.0028, two-tailed unpaired t-test, **Figure 6b**) higher than those injected with vehicle at 48 h after activation, while we did not observe significant difference at 24 h time point (P=0.227, two-tailed unpaired t-test, **Extended data Fig. 9a**). We next found that mice injected with CNO generated plasma Nluc-RMA signals, which were derived from *Arc* sensing at the striatum, that were 1.98 ± 0.53-fold higher than those injected with vehicle at 48 h after activation (P=0.0048, two-tailed unpaired t-test, **Figure 6c**). Additionally, we found that Nluc-RMA signal was also significantly higher at 24 h after the CNO injection, as compared to the vehicle (1.67 ± 0.42-fold, P=0.0111, two-tailed unpaired t-test, **Extended data Fig. 9b**). We reasoned that since Nluc-RMA outperformed the sensitivity over Gluc-RMA, Nluc-RMA successfully captured the lower induction of *Arc* expression in the striatum, where Gluc-RMA failed (**Extended data Fig. 6h, i**). As expected, the negative control Cluc-RMA signals, driven by hSYN promoter in the midbrain, were unchanged by chemogenetic induction (P=0.9532, and P=0.6374 at 48 h and 24 h time points, respectively, two-tailed unpaired t-test, **Figure 6d**, and **Extended data Fig. 9c**, respectively). After 48 h, we perfused the animals to analyze the brain tissue histologically. We found that the number of cells expressing Gluc-RMA, Nluc-RMA, and Cluc-RMA were 1.83 ± 0.19-fold, 2.37 ± 0.68-fold, and 0.99 ± 0.65-fold higher in the CNO group than in the vehicle group (P<0.0001, P=0.0059, NS P>0.05 respectively, two-tailed unpaired t-test, **Figure 6e–h, Supplementary Fig. 7–9**), which was consistent with blood RMA signals.

## DISCUSSION

We established INTACT as a new paradigm for non-invasive measurement of sequence-specific mRNA levels in the living brain with a simple blood test. INTACT relies on a combination of synthetic serum markers and RNA-based sensors of mRNA. The synthetic markers are expressed in the brain, but exit into the blood where they can be easily measured. The sensors tie that expression to the levels of sequence-targeted mRNA within the cell. Here, we demonstrate that INTACT can measure mRNA *in vivo*. INTACT showed high signal levels with up to 1,294-fold RMA signal over the baseline, depending on the type of the probe used and the detected transcript. Post-mortem histology showed that the specificity of RMA expression was consistent between the tested sensors. For the detection of tdtomato, *TH, Vgat, c-Fos*, and *Arc* transcripts, we observed up to 94.1% of cells that expressed RMAs *in vivo*, also expressed proteins translated from these transcripts. Interestingly, INTACT readout matched the previously-reported patterns of gene expression in the brain. For example, c-Fos expression was elevated in the striatum after chemogenetic activation^17,40,^ while Arc was activated in the CA1 of the hippocampus more strongly than the striatum under physiological condtions^41^.

INTACT holds several advantages over existing techniques for monitoring gene expression. Compared to tissue-destructive methods, such as qPCR, INTACT can monitor the history of gene expression over time. Compared to existing noninvasive imaging methods, such as protein-based MRI contrast agents^14,42-45,^ biomolecular ultrasound^13,46,^ or BLI, INTACT provides simpler and more multiplexable interface for readout. For example, while BLI can provide readout of several colors of luciferases in one sample, detection of proteins in the serum can be done through massively multiplexed methods, such as mass spectrometry^47^ or single-protein sequencing^48,49,^ to provide noninvasive measurement of transcriptomic information. Thirdly, unlike in biomolecular ultrasound or BLI, INTACT is not affected by the tissue thickness, skull presence, or tissue opacity, resulting in comparably higher signal levels. For example, in our previous studies *Gaussia* luciferase expressed in the mouse brain, and imaged through intravital imaging, showed over 100-fold lower signal, than when it was released into the blood as an RMA^17^. Lastly, compared to these methods INTACT readout is more flexible and simpler without needing immobilization of animals in expensive scanners. The INTACT concept thus democratizes access to the deep-tissue gene expression monitoring, while improving the sensitivity, and providing a higher theoretical ceiling on the multiplexity of readout.

Recent discovery of “ticker tapes” that can record gene expression history in individual cells is a groundbreaking development^50,51^. These technologies use an ordered multi-protein structure that is assembled in a pattern dependent on the level of reporter protein expression at a given time. ‘Ticker tapes’ are particularly well suited for monitoring gene expression histories in individual cells in known parts of the tissues. A similar technology, called peCHYRON recently has been developed to record cellular events, such as gene expression patterns in the cell’s genome^52^, which can be recorded with DNA sequencing. However, the readout of ‘ticker tapes’ or peCHYRON, is currently done either in post-mortem tissues, or cell culture. INTACT on the other hand, reads the gene expression histories nondestructively and is thus ideal for applications where the animal is kept alive and the tissue needs to remain intact, such as in large animal, long-term, or potential clinical studies. Another interesting technology uses engineered retroviral capsids to secrete intracellular RNA. However, this technology so far has not been adapted for *in vivo* use^53^.

As shown here, INTACT records from the genetically-labeled cell populations, with the spatial precision being defined by the region of gene delivery of the INTACT’s components. However, our recent work has shown that RMAs can be read-out in intact tissues with millimeter spatial precision through the use of focused-ultrasound (FUS) blood-brain barrier opening (FUS-BBBO)^54^.

One potential limitation of ADAR-mediated RNA sensors is their efficiency. For example, ADAR-mediated RNA editing was observed in only 3% of cultured cells receiving similar sensor and target constructs^55^. To date, the RNA-targeting sensors detected endogenous mRNA *in vivo* with invasive imaging methods and post-mortem^21^, but their application to detection through noninvasive monitoring of transcripts could be challenging. Despite the potential limited efficacy of ADAR, we observed detectable levels of RMA expression driven by ADAR-based sensors, with signals comparable to floxed RMAs expressed under hSyn1 promoter^17^. The signal sensitivity and target specificity of the INTACT platform could be further improved by optimizing the target and length of sensor binding sequences, varying the number of stop codons, overexpressing ADAR enzymes, incorporating MS2 hairpin loops to recruit ADAR proteins^56,57^, and circularizing RNA sensor^58^. Since RNA sensor technologies are a recent breakthrough^21-24,59-61^, new advancements in this field are expected and will also benefit INTACT concept.

Another limitation of INTACT platform is common with other genetically-encoded reporters. Specifically, INTACT also requires gene delivery. Here, we used invasive delivery through intracranial injection of AAVs carrying INTACT constructs, while making the readout minimally invasive through blood draws. This gene delivery approach is commonly used across the neuroscience laboratories owing due to its simplicity, spatial precision, and low amount of the vector needed. However, other minimally-invasive gene delivery methods exist, including BBB-permeable AAVs^62^ that can target specific cell populations through cell-type specific sensors, or focused ultrasound blood-brain barrier opening (FUS-BBBO) delivery of AAVs^63-65^ that can additionally provide spatial precision. However, with a systemic delivery to the brain, care needs to be taken to distinguish any residual expression from peripheral tissues, a limitation which we addressed in our recent study^54^.

In the current study, we focused on using luciferases as synthetic serum markers due to their simple detection. However, in the next steps, integration of peptide barcodes for single-molecule protein sequencing^48,66^, or single-chain variable fragments of antibodies (scFv) would enable highly multiplexed readout. A second area of improvement could include reduction of off-target editing of the non-target adenine within the synthetic RNA sensor^21-23^. Further engineering ADAR could maximize the target specificity of editing. We previously demonstrated no detectable immunogenicity of RMA reporters in the brain and peripheral organs^17^ until at least 1,000-fold signal over the baseline. However, further investigation will be needed to evaluate long-term safety of various types of RMAs. Lastly, development and testing of INTACT in other species will be valuable. In particular, application of INTACT in large animal species, such as non-human primates, will be transformative, as post-mortem brain tissue collection in these species poses substantial practical and ethical challenges, and noninvasive methods of monitoring gene expression, such as bioluminescence imaging, are hampered by the signal attenuation through a thick skull and the overall large size of the brain. Collectively, we anticipate that these improvements can transform the INTACT platform for highly precise, noninvasive monitoring of gene expression, facilitating biological discoveries and developing next-generation precision diagnostics and therapeutic strategies^67^.

## MATERIALS AND METHODS

### Animal subjects

WT C57BL/6J (strain 000664) were purchased from The Jackson Laboratory and used for experiments at 9–12 weeks of age. Animals were housed with a 12-h light/dark cycle, 18– 23 °C ambient temperature and 40–60% humidity and were provided with food and water ad libitum. All animal experiments were performed under the protocol approved by the Institutional Animal Care and Use Committee of Rice University.

### Plasmid construction

All constructs were generated using standard cloning procedures. Vector backbones were linearized using restriction digestion, and DNA fragment inserts were generated using PCR or gBlock synthesis (Integrated DNA Technologies). To construct CAG-BFP-sesRNA-Gluc-RMA, the vector CAG-BFP-sesRNA-GFP (Addgene, 192063) was digested with PacI and HindIII (New England Biolabs) to isolate the backbone. Gluc-RMA was amplified by polymerase chain reaction (PCR) from the vector AAV-hSyn1-Gluc-RMA-IRES-EGFP (Addgene, 189629), and its DNA was extracted using a Zymoclean Gel DNA Recovery Kits (ZYMO). Gluc-RMA was inserted into the digested backbone through Gibson assembly. To construct AAV-hSyn1-EGFP-sesRNA-tTA2, the BFP in the vector CAG-BFP-sesRNA-tTA2 (Addgene, 192070) was replaced with EGFP. To construct TRE3g-Gluc-RMA, the vector AAV-TRE3g-mNeon (Addgene, 192064) was digested with SalI and SpeI (New England Biolabs) to isolate the backbone. Gluc-RMA was inserted into the digested backbone as above. To construct TRE3g-Nluc-RMA, the Gluc region of the vector TRE3g-Gluc-RMA was replaced by the sequence of secreted-type nanoluc luciferase (GenBank: JQ437372.1). To construct CAG-ADAR2-sesRNA-RMA, the vector CAG-ADAR2-sesRNA-GFP (Addgene, 192069) was digested with PacI and NheI (New England Biolabs) to isolate the backbone. Gluc-RMA was inserted into the digested backbone as above.

To construct CAG-TH and CAG-Arc, the vector CAG-tdTomato (Addgene, 83029) was digested with EcoRI and BglII to isolate the backbone. TH sequence was cloned from the pISH-TH (Addgene, 105988) into the backbone. *Arc* coding sequence was cloned from the mouse cDNA library into the backbone. All plasmid information and sensor sequences (sesRNA) used in this study are included in **Supplementary Table**.

### Sensor RNA design

For sesRNA design, we followed the procedures previously reported^21^. Briefly, sesRNA – a complementary sequence to the target mRNA sequence, satisfies the following requirements: (1) length of sesRNA is 150-400bp; (2) one or more stop codons (TAG) are placed near the center of a sesRNA. This stop codon creates an A-C mismatch that could be modified by ADAR. (3) ensure that there are no other stop codons in the sesRNA, all other TAG, TAA, and TGA sequences in sesRNA are converted to GAG, GAA, and GGA, respectively. (4) no initiation codon (ATG) is recommended after the TAG in sesRNA to decrease the possibility of unintended translation initiation. (5) avoid sesRNA with complex secondary structures. For targeting tdTomato and *Vgat* sequences, their sesRNA were referred to the previously established sesRNA^21^ respectively.

### Cell culture for luciferase assay

PC-12 (American Type Culture Collection, CRL-1721) was cultured in RPMI 1640 medium (Corning, 10-040-CV) supplemented with 10% horse serum (Life Technologies, 26-050-088) and 5% FBS (Corning, 35-011-CV). HEK293T (American Type Culture Collection, CRL-3216) was cultured in Dulbecco’s Modified Eagle Medium (DMEM) (Corning, 10-013-CV) supplemented with 10% FBS. Cells were incubated in humidified air with 5% CO2 at 37 °C and split every 2 d with a subcultivation ratio of 1:2 or 1:3.

For in vitro luciferase assay, PC-12 or HEK293T were seeded at 100,000 cells per well in a 24-well plate. After 16–20 h, 250 ng of plasmids encoding RNA sensor, RMA reporter, mRNA target, exogenous ADAR, and 2.5 µl of Lipofectamine 2000 (Life Technologies, 11-668-019) were used for transfection, following the manufacturer’s protocol. Then, 25 µl of the culture media was collected at different time points and stored at −20 ^°^C. For Gluc substrate, 0.5 mM native coelenterazine (CTZ) stock (Nanolight Technology, 303) was dissolved in luciferase assay buffer (10 mM Tris, 1 mM EDTA, 1.2 M NaCl, pH 8.0) containing 66% DMSO and stored at −80 ^°^C. Before measuring bioluminescence, the CTZ stock was diluted to 20 µM in luciferase assay buffer and kept in dark at room temperature for 1 h. Media samples were thawed and transferred to a black 96-well plate (Corning). Infinite M Plex microplate reader equipped with i-control software (Tecan) was used to inject 50 µl of the assay buffer containing CTZ and measure the photon emission integrated over 30 s. For measuring bioluminescence of Cluc or Nluc, samples were mixed by injecting 50 µl of 1.0 µM vargulin (Nanolight Technology, 305), or 100 times diluted Nluc substrate (Promega, N2011) respectively, dissolved in the luciferase assay buffer. Values were averaged to obtain the light unit per second using Excel (Microsoft).

### RT-qPCR

RNA was extracted and purified with RNeasy Mini Kit (Qiagen, 74106). RNA was converted to cDNA using Verso cDNA Synthesis kit (Thermo Fisher, AB1453B). RT-qPCR was performed with optimal primer sets and PowerUp™ SYBR™ Green Master Mix for qPCR (Thermo Fisher, A25741) on the qTOWER 3G Real-Time PCR Thermal Cycler (Analytik Jena). The primers were synthesized from Sigma (18S; Forward: ACCGCAGCTAGGAATAATGGA, Reverse: GCCTCAGTTCCGAAAACCA, tdTomato; Forward: ATCGTGGAACAGTACGAGCG, Reverse: TGAACTCTTTGATGACGGCCA).

### AAV production

PHP.eB AAV virus was packaged by adopting a previously published protocol with slight modifications. In brief, HEK293T cells (American Type Culture Collection, CRL-3216) were transfected with the transfer plasmid, PHP.eB iCap (Addgene, 103005) and pHelper plasmids. After 24 hours, cells were exchanged with a fresh DMEM (Corning, 10-013-CV) supplemented with 5% FBS and non-essential amino acids (Life Technologies, 11140050). At 4 days after transfection, cells were harvested, and medium were mixed with 1/5 volume of PEG solution (40% PEG 8,000, 2.5 M NaCl) to precipitate AAV at 4 ^°^C for 2 hours. Cells were resuspended in PBS and lysed by the freeze-thaw method. The precipitated AAV was pelleted by centrifugation, resuspended in PBS and combined with the lysed cells. The combined lysate was added with 50 Uml-1 Benzonase (Sigma-Aldrich, E1014) and incubated at 37 ^°^C for 45 min.

AAV purification was carried out by the iodixanol gradient ultracentrifugation. A Quick-Seal tube (Beckman Coulter, 344326) was loaded with the iodixanol gradients (Sigma-Aldrich, D1556), including 60%, 40%, 25% and 15%. The lysate was centrifugated, and the resulting clarified lysate was loaded on the top of the iodixanol layers. The tube was sealed and centrifugated at 350,000g for 2.5 hour using a Type 70Ti fixed-angle rotor of an ultracentrifuge (Beckman Coulter). AAV was collected by extracting the 40–60% iodixanol interface and washed using the Amicon centrifugal filter unit with 100-kDa cutoff (MilliporeSigma). The final AAV was filtered by passing through the 0.22-µm PES membrane. Viral titers were determined using the qPCR method.

### Stereotaxic injection

AAV was injected into the mice brains using a microliter syringe equipped with a 34-gauge beveled needle (Hamilton) installed to a motorized pump (World Precision Instruments) using stereotaxic frame (Kopf Instruments). For AAV injections, PHP.eB serotype was used for all experiments. For tdTomato targeting, 0.5 x 10^9^ vg of each AAV encoding hSyn1-EGFP-sensor (tdT)-tTA2, TRE-RMA, CAG-tdTomato, and hSyn1-Cluc-RMA are mixed in 200 nl and injected per site at 600 nl min^-1^ to the following coordinates: CP in the striatum (AP +0.25 mm, ML + 2.0 mm, DV −3.2 mm). For TH targeting, 0.5 x 10^9^ vg of each AAV encoding hSyn1-EGFP-sensor (TH)-tTA2, TRE-RMA, and hSyn1-Cluc-RMA are mixed in 200 nl and injected per site at 600 nl min^-1^ to the following coordinates: VTA (AP -2.9 mm, ML + 0.8 mm, DV −4.55 mm). For *Vgat* targeting, 0.5 x 10^9^ vg of each AAV encoding hSyn1-EGFP-sensor (Vgat)-tTA2, TRE-RMA, and hSyn1-Cluc-RMA are mixed in 200 nl and injected per site at 600 nl min^-1^ to CP in the striatum.

For chemogenetic neuromodulation, mice were placed on 40 mg kg^−1^ Dox chow (Bio-Serv) 24 h before surgery. AAV doses used are as follows: 0.5 × 10^9^ vg (hSyn1-EGFP-sensor (c-Fos or Arc)-tTA2), 0.5 × 10^9^ vg (TRE3g-Gluc-RMA), 0.5 × 10^9^ vg (hSyn1-hM3Dq-mCherry) and 0.5 × 10^9^ vg (hSyn1-Cluc-RMA). For each surgery, AAV cocktail was prepared in 200 nl and injected per site at 600 nl min^-1^ to CP in the striatum or CA1 in the hippocampus (AP -1.94 mm, ML + 1.0 mm, DV −1.45 mm). Dox chow was removed 48 h before inducing chemogenetic activation. For triplex monitoring, AAV doses were used are as follows: 0.5 × 10^9^ vg (hSyn1-EGFP-sensor (Arc)-tTA2), 0.5 × 10^9^ vg (TRE3g-(Gluc or Nluc)-RMA), 0.5 × 10^9^ vg (hSyn1-hM3Dq-mCherry) and 0.5 × 10^9^ vg (hSyn1-Cluc-RMA). The AAV encoding hSyn1-EGFP-sensor (Arc)-tTA2 was injected into the CA1 in the hippocampus with AAV encoding TRE3g-Gluc-RMA. The AAV encoding hSyn1-EGFP-sensor (Arc)-tTA2 was also injected into the striatum (AP +0.25 mm, ML −2.0 mm, DV −3.2 mm) with AAV encoding TRE3g-Nluc-RMA. AAV encoding hSyn1-Cluc-RMA was injected into the SN in the midbrain (AP −3.28 mm, ML +1.5 mm, DV −4.3 mm).

### Blood collection for luciferase assay

Mice were anesthetized in 1.5–2% isoflurane in air or O2. Next, 1–2 drops of 0.5% ophthalmic proparacaine were applied topically to the cornea of an eye. A heparin-coated microhematocrit capillary tube (Thermo Fisher Scientific, 22-362566) was placed into the medial canthus of the eye, and the retro-orbital plexus was punctured to withdraw 50–100 µl of blood. The collected blood was centrifuged at 1,500g for 5 min to isolate plasma. To conduct luciferase assay, 5 µl of plasma was mixed with 45 µl of PBS + 0.001% Tween 20 in a black 96-well plate. Using the microplate reader, bioluminescence of Gluc-RMA, Cluc-RMA, or Nluc-RMA was measured by injecting 50 µl of 20 µM CTZ, 1.0 µM vargulin, or 100-times diluted Nluc substrate, respectively, dissolved in the luciferase assay buffer.

### Drug administration

Water-soluble CNO (Hello Bio, HB6149) was dissolved in saline (Hospira) at 1 mg ml^−1^. To induce chemogenetic activation of mice expressing hM3Dq, CNO was injected intraperitoneally at 2.5 mg kg^−1^ dose. For the vehicle (0 mg kg^−1^) groups, saline was injected intraperitoneally at the volume dose of 5 ml kg^−1^.

### Immunohistochemistry

Mice brains were extracted and post-fixed in 10% neutral-buffered formalin (Sigma-Aldrich, HT501128). Coronal sections were cut at a thickness of 50 µm using a vibratome (Leica). Sections were stained as follows: (1) block for 1 h at room temperature with blocking buffer (0.2% Triton X-100 and 10% normal donkey serum in PBS); (2) incubate with primary antibody overnight at 4 ^°^C; (3) wash in PBS for 15 min three times; and (4) incubate with secondary antibody for 4 h at room temperature. After the last washes in PBS, sections were mounted on glass slides using the mounting medium (Vector Laboratories). Antibodies and dilutions used are as follows: rabbit anti-Gluc (1:1,500, Nanolight Technology), mouse IgG2a anti-Fos (1:500, Santa Cruz Biotechnology), Guinea pig anti-Arc (1:500, Synaptic Systems), mouse IgG2a anti-Nanoluc (1:500, Promega) and Alexa 350, 488, 594 and 647 secondary antibodies (1:500, Life Technologies).

All images were acquired by a BZ-X800 fluorescence microscope (Keyence). Manual cell counting was performed using ZEN Blue software (Zeiss). Cells in all images were counted individually. Cell specificity of sensor RNA was calculated as the ratio of positive cells expressing target protein or mRNA among the cells expressing the RMA reporters. Sensor efficiency was calculated as the ratio of RMA-positive cells over RNA sensor (GFP) and target protein (tdTomato). The manual cell counts were validated by a blinded observer and determined to be within 7.4% on average and specifically: for Fig. 3d, 12.1% (vehicle, P=0.109), 8.3% (CNO, P=0.096) specificity for Fig. 5h, 13.0% (P=0.645) of fold change for Fig. 5g, 5.32% (P=0.614) of fold change for Fig. 6f, 13.9% (P=0.116) of fold change for Fig. 6g, 1.14% (P=0.932) of fold change for Fig. 6h, 14.2% (P=0.352) of fold change for Extended data Fig. 5d, 6.82% (P=0.824) of fold change for Extended data Fig. 5j. To score the RMA localization, representative images of nuclear, both of nuclear and peripheral, or primarily peripheral localization were defined as the score 0,1, and 2, respectively. Then, other images were manually scored by blinded observer uninformed of the experimental conditions.

For the cell counts in the tissue, the values reported refer to the targeted brain region, as defined by the area of expression of the INTACT components within the brain.

### In situ hybridization

All probes were ordered from Molecular Instruments. Mouse brain slices were prepared as described in immunohistochemistry. The hybridization chain reaction in situ was performed via free-floating method in a 24-well plate. First, Brain slices were exposed to probe hybridization buffer with HCR probe Set (Molecular Instruments Inc.) at 37°C for 24 h. Brain slices were washed with probe wash buffer, incubated with amplification buffer, and amplified at 25°C for 24 h. On day 3, brain slices were washed, counter stained using immunohistochemistry.

### Statistical analysis

Two-tailed t-test with unequal variance was used to compare two datasets. One-way ANOVA with Tukey’s or Dunnett’s honestly significant difference post hoc test was used to compare means between more than two datasets. Two-way ANOVA with Sidak’s multiple comparison tests was used to compare datasets with two or more variables. Michaelis-Menten analysis was used to find the correlation between the plasma luciferase signal and the tdTomato mRNA expression. All P values were determined using Prism (GraphPad Software), with the statistical significance represented as not significant (NS), P ≥ 0.05; *P < 0.05; **P < 0.01; ***P < 0.001; and ****P < 0.0001. P values and statistical test results are available in the Source Data. Figures were constructed using Adobe Illustrator. All numerical values are provided as means with ± standard deviations.

## Data availability

The authors declare that all data supporting the results in this study are available within the paper, its Supplementary Information and its Source Data file. Microscopy images are available from the corresponding author upon reasonable request owing to their large size and number. Plasmids available upon request. Source data are provided with this paper.

## ACKNOWLEDGEMENTS

We thank Adam E. Cohen (Harvard University) for helpful suggestions during preparation of the manuscript. This research was supported by the David and Lucile Packard Foundation 2021-73005 to J.O.S., National Institute of Biomedical Imaging and Bioengineering award DP2EB035905 to J.O.S, and Japan Society for the Promotion of Science fellowship to S.W.

## AUTHOR CONTRIBUTIONS

J.O.S. and S.W. conceived and planned the research. S.W. and J.O.S. designed the experiments and wrote the paper, with input from all other authors. S.W. performed and participated in all experiments described in the study. S.L. developed Gluc-RMA and Cluc-RMA. S.L. confirmed the orthogonality between three different RMAs in vitro. S.N., E.R. and H.L. performed the histological experiments. M.H, E.R. and H.L. analyzed histological images. M.H. and N.B. conducted the quantification of intracellular RMA localization. B.P. developed Nluc-RMA and confirmed its function in vitro.

## Competing interests

The authors declare no competing interests.

## Notes

### Competing Interest Statement

The authors have declared no competing interest.

### Summary of Updates

adding extended data figures to the supplementary file

