## Extended data figures for "Monitoring in vivo transcription with synthetic serum markers"

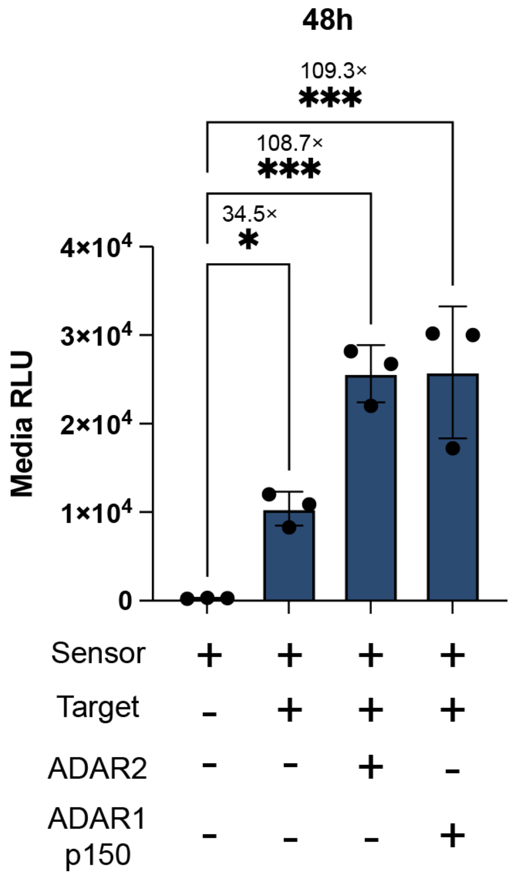


**Extended Data Figure 1. The effects of exogenous ADAR expression on RMA output**
RMA secretion in response to tdTomato expression with and without ADAR overexpression. RLU values measured from the culture media reveal the target mRNA-dependent secretion of RMA. ADAR2 was expressed from the same plasmid as the RNA sensor. ADAR1p150 was expressed in trans from a separate plasmid. Plasmid details are described in Supplementary Table. n=3 independent cultures were analyzed. In comparison with the signal in the group of no-target, using one-way ANOVA ($F_{3,8}$=26.88, P=0.0002), with Dunnett's test: P=0.0435 (no-target versus Target), P=0.0002 (no-target versus Target with ADAR2), and P=0.0002 (no-target versus Target with ADAR1p150). *P<0.05 and ***P<0.001. Data is shown as mean ± s.d.

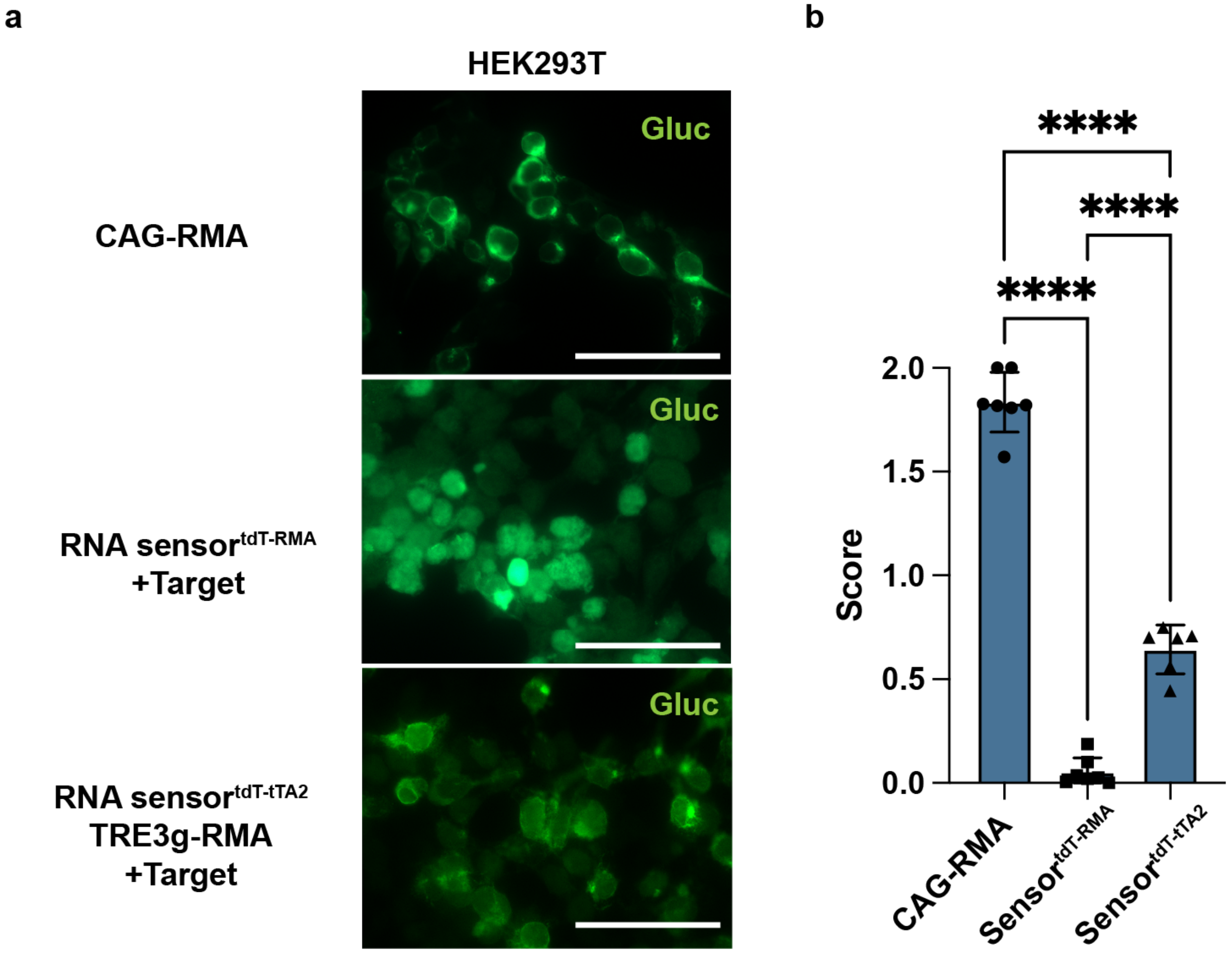


**Extended Data Figure 2. Intracellular localization of RMA protein between different gene circuits**

**(a)** Immunostaining images of the HEK293T cells transfected by CAG-RMA, tdTomato-targeting sensor (**Figure 2a**), and tTA2-TRE-based sensor (**Figure 2d**). RNA sensors were co-transfected with CAG-tdT into PC12 cells as a target mRNA to drive RMA expression. Intracellular Gluc-RMA localization was visualized through immunostaining with Gluc antibody. Scale bars, 50 μm. **(b)** The localization patterns of RMA protein were manually quantified by the blinded observer. The representative images of each group are scored 2 for primarily peripheral location of RMA, 1 for presence of visible peripheral localization along with the cytoplasmic signal, and 0 with indistinguishable peripheral signal. The mean score was calculated as the final score in the image. n=7 randomly selected images from N=2 independent cultures were used for analysis. P<0.0001 for all comparisons, using one-way ANOVA ($F_{2,17}$, P=441.5) with Tukey's test. ****P<0.0001. Data are shown as mean ± s.d.

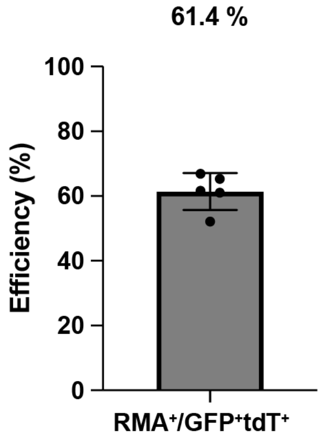


**Extended Data Figure 3. Efficiency of tdT-targeting RNA sensor *in vivo***
Efficiency of sensing is calculated as the ratio of RMA positive cells among the cells that also express the sensor and tdT target. Data are shown as mean ± s.d.

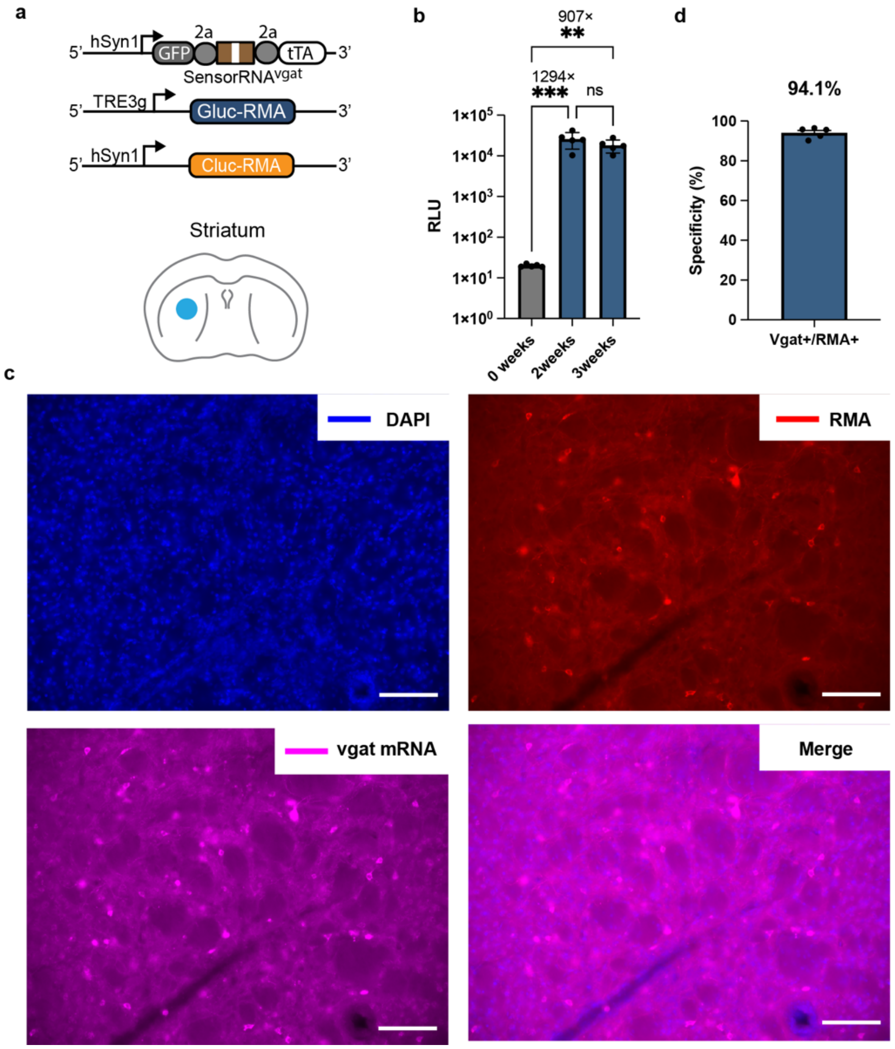


**Extended Data Figure 4. Monitoring inhibitory neurons with *Vgat* mRNA sensor.**

**(a)** Experimental design for detection of *Vgat* mRNA *in vivo*. Three different AAVs, each at $0.9\times10^{9}$ vg dose were injected into the mouse striatum. The AAVs encoded vgat-targeting RNA sensor, TRE-Gluc-RMA reporter, and hSYN1-Cluc-RMA control. (b) Afterwards, blood was collected to measure RMA levels. Signals were increase at both two (P=0.0004) and three weeks (P=0.0065) time points as compared to the 0-week baseline. n=5 independent mice were analyzed using one-way ANOVA ($F_{2,12}$=15.55, P=0.0005) with Tukey's test. **p<0.01, ***p<0.001. **(c)** Representative images showing expression of Gluc-RMA (red) and *Vgat* mRNA (magenta). Gluc-RMA was labeled by Gluc antibody, while *Bgat* mRNA was labeled by in situ hybridization. Scale bars 100 μm of the enlarged images. **(d)** The specificity of *Vgat* targeting by the RNA sensor. Specificity was assessed by counting the number of RMA-positive cells that were also labeled with *Vgat* mRNA. n=5 The data from independent mice were analyzed. Data is shown as mean ± s.d.

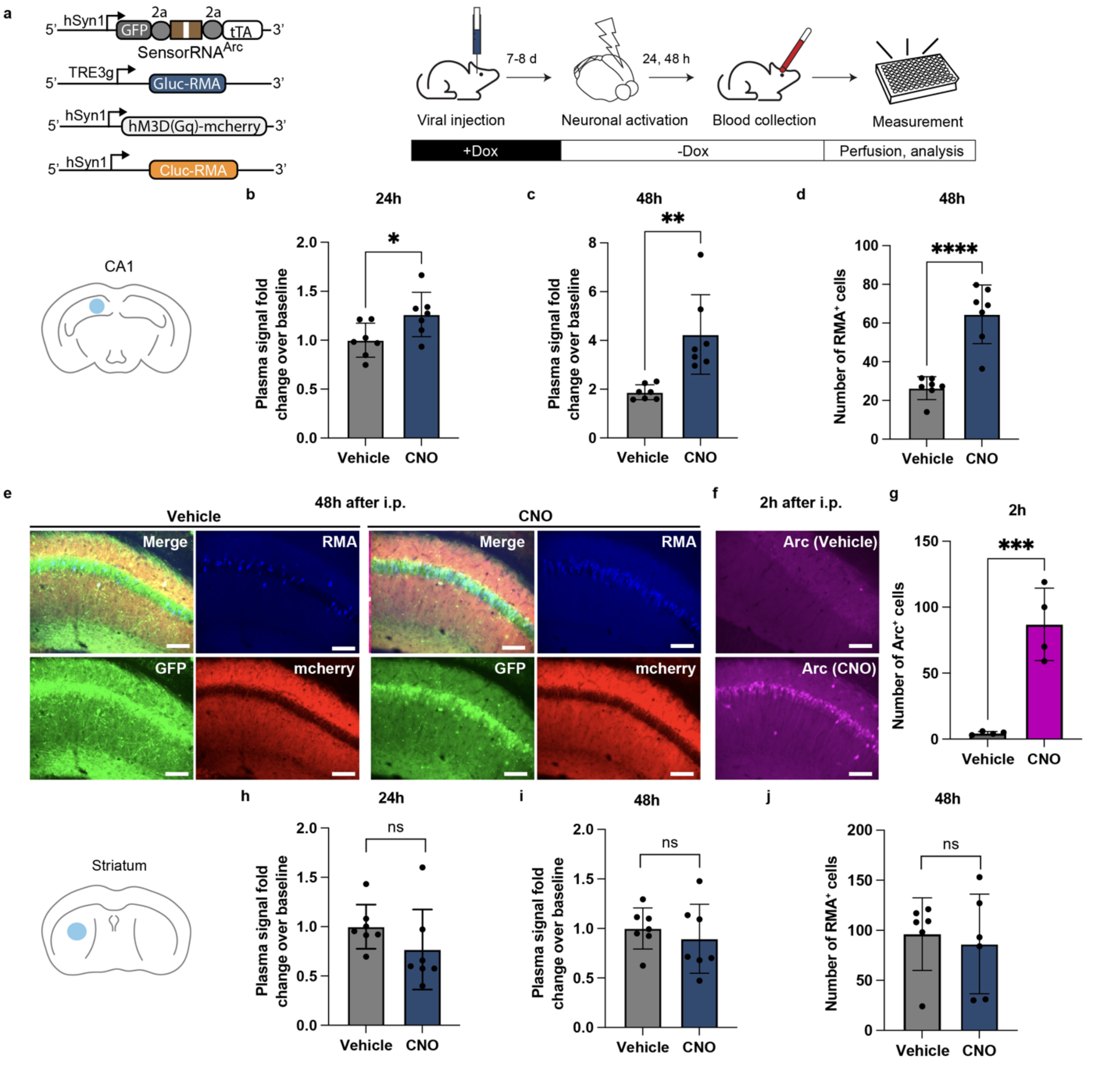


**Extended Data Figure 5. Measurement of *Arc* transcripton in the brain with INTACT.**

**(a)** Experimental scheme for detection of *Arc* mRNA with INTACT in response to chemogenetic neuronal activation. AAVs encoding an *Arc* RNA sensor (Syn-sesRNA (Arc)-tTA2), TRE-Gluc-RMA, activatory DREADD (hM3Dq) receptor, and hSyn1-Cluc-RMA were injected into the brain. Mice were then placed on the Dox diet to shut down the RMA production until the desired timepoint for *Arc* recording. After 7 days, the Dox diet was withdrawn, and 48 hours later mice received either CNO or vehicle injections. Blood was then drawn at 24h and 48h to analyze the serum RMA levels. **(b)** RMA signal fold change as comparted to the 0 h baseline after chemogenetic stimulation in the hippocampus CA1 at 24h, or **(c)** 48 h after CNO or vehicle injection. Signals were normalized by dividing the RLU values obtained from Gluc-RMA over

Cluc-RMA. n=7 (vehicle) and n=7 (CNO) independent mice were analyzed using unpaired two-tailed t-test at 24 h (P=0.0317) and 48 h (P=0.0026) time points. *P<0.05, **P<0.01. **(d)** The number of RMA-expressing cells at 48 h CNO or vehicle injection (n=7 mice per group). P<0.0001 using unpaired two-tailed t-test. ****P<0.0001. **(e)** Representative images showing expression of the RNA sensor (GFP), DREADD Gq receptor (mCherry), and Gluc-RMA (Blue) in the targeted brain region at 48 h after CNO or vehicle administration. Scale bar, 500 μm. **(f)** Representative images of *Arc* protein expression (magenta) in the mouse CA1 at 2 h after CNO or vehicle injection. Scale bar, 500 μm. **(g)** The number of Arc-expressing cells in the targeted brain region at 2 h after CNO or vehicle administration (n=4 mice per group). P=0.0010 using unpaired two-tailed t-test. ***P<0.001. **(h)** RMA signal fold change as comparted to the 0 h baseline after chemogenetic stimulation in the striatum at 24h or **(i)** 48 h. Signals were normalized by dividing the RLU values obtained from Gluc-RMA over Cluc-RMA. N=7 mice were analyzed in each group using unpaired two-tailed  t-test at 24 h (P=0.2106 (b) and 48 h (P=0.5105) (c) time points. ns P>0.05. **(j)** The number of Arc-expressing cells in the targeted brain region. n=7 (vehicle) and n=6 (CNO) independent mice were analyzed. P=0.7038 using unpaired two-tailed Student's t-test. ns P>0.05. Data is shown as mean ± s.d.

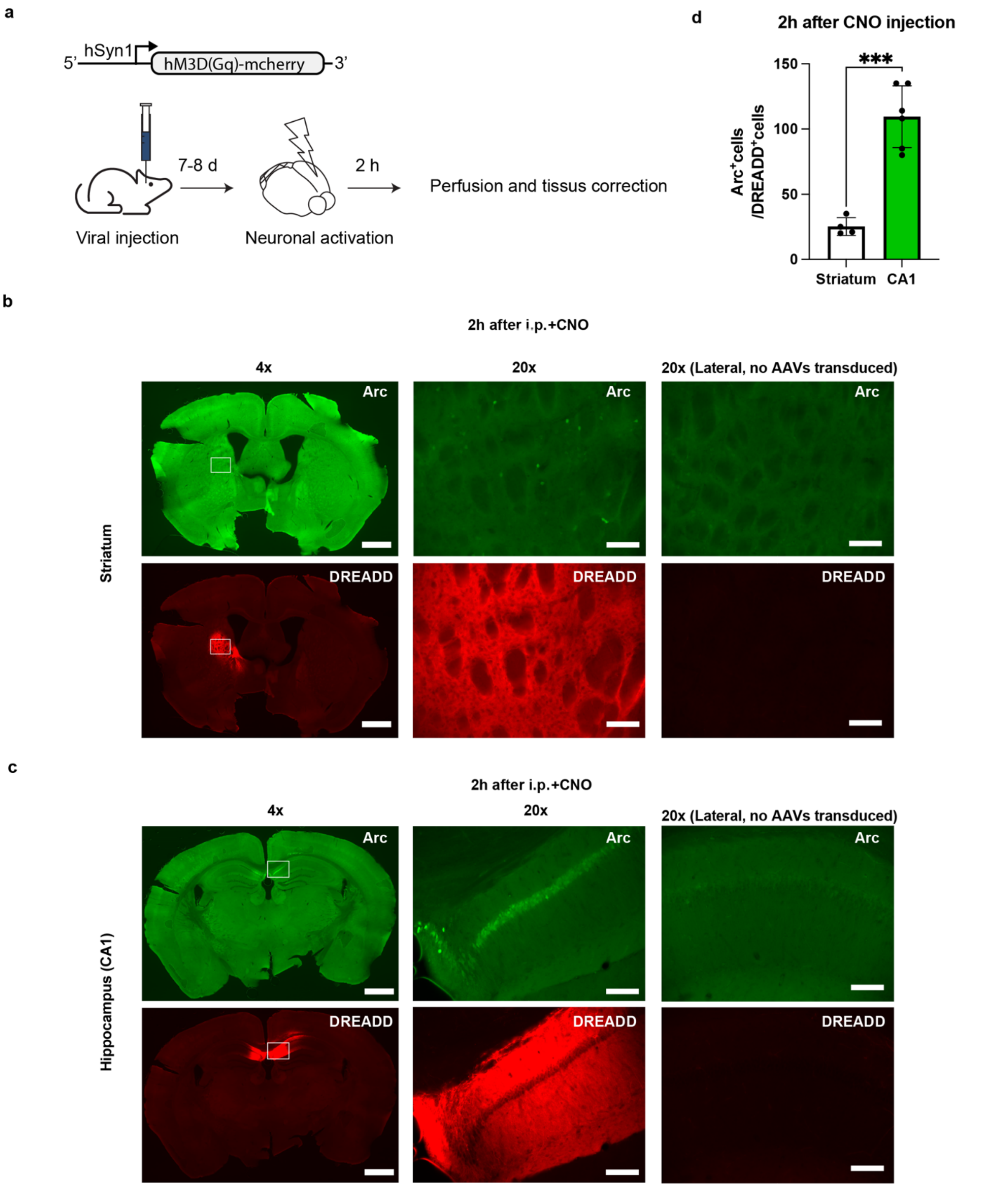


**Extended Data Fig. 6. Region-specific induction of *Arc* expression by DREADD-induced neuronal activation.**

(a) Experimental scheme for detection of *Arc* protein expression in response to DREADD-induced neuronal activation. Mice undergo injection of AAV encoding activator DREADD (hM3Dq) receptor, followed by by CNO or vehicle injection. 2 h after intraperitoneal injection (i.p.) injection of CNO, brain tissue was perfused and collected for histology analysis. **(b, c)** Representative images of protein expression of *Arc* (Green) and DREADD Gq (mCherry) at striatum (b) and hippocampus CA1 (c). The tissue collected 2 h after i.p. from the CNO-injected mice. Scale bars 1000 μm and 100 μm of the left (whole brain) and the middle and right (enlarged) images, respectively. The right images are the lateral sides of brain tissue where no AAVs were transduced. **(d)** Propensity of *Arc* to be induced by DREADD activation as defined by the number of Arc-positive cells divided by the number DREADD-expressing cells at 2 h after CNO or vehicle injection n=4 (striatum) and n=6 (CA1) brain slices analyzed from from N=2 independent mice. P=0.0001 using unpaired two-tailed t-test. ***P<0.001. Data is shown as mean ± s.d.

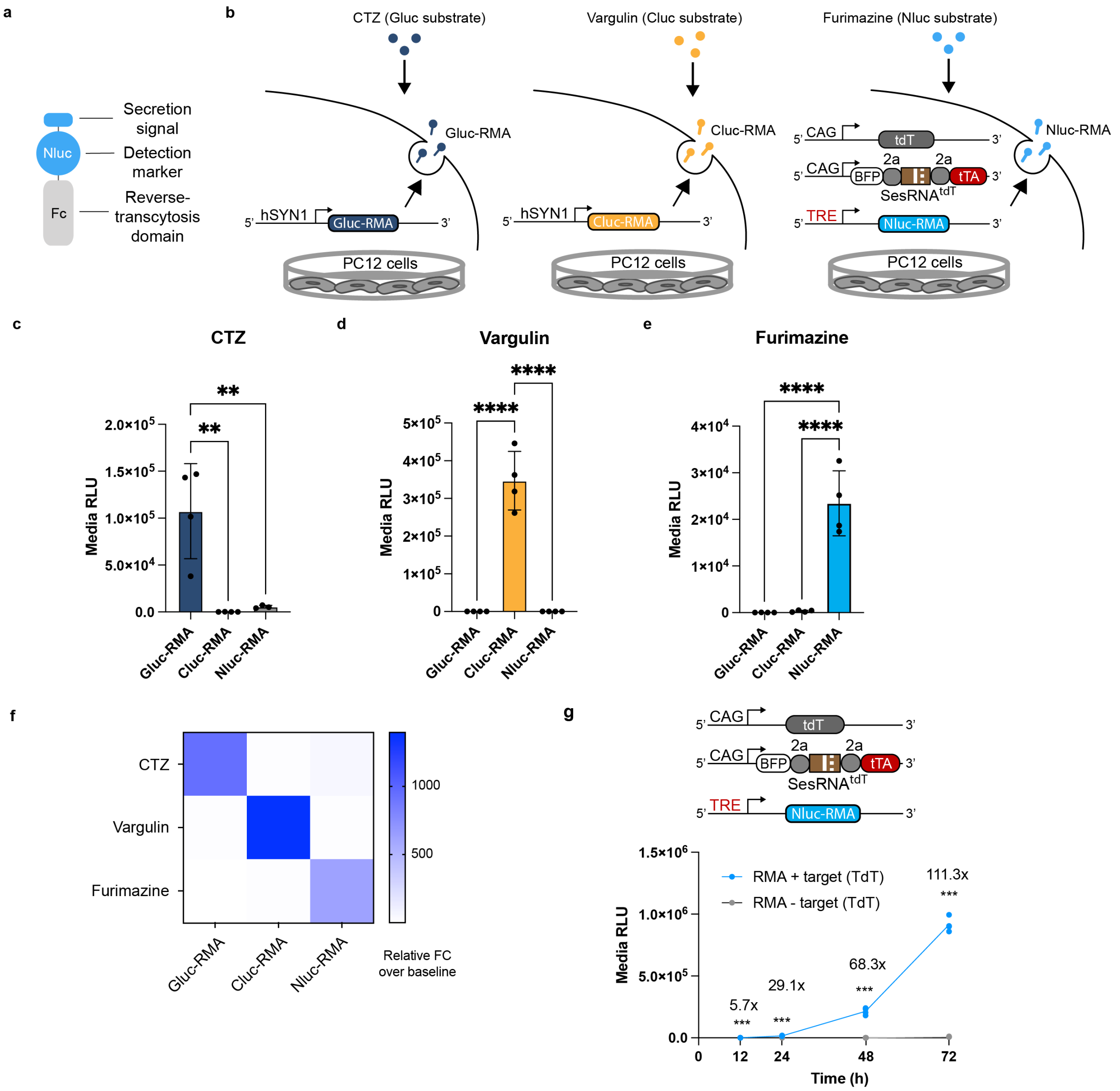


**Extended Data Fig. 7. Development of Nluc-RMA reporter and evaluation of functional orthogonality between three different RMAs.**

**(a)** Nluc-RMAs design include a secretion signal along with the Fc region of the IgG2 antibody. The sequence information for Nluc was referred to GenBank (JQ437372.1). **(b)** Experimental design for testing the orthogonality of detection for each RMA. Gluc-RMA and Cluc-RMA were driven by hSYN promoter while Nluc-RMA, was driven by an RNA sensor circuit targeting tdT. CAG-tdT encoded tdTomato providing target mRNA. The culture medium was collected at 48 h after transfection to measure the luminescence activity. CTZ, vargulin, and furimazine were used for detecting Gluc, Cluc, and Nluc activity, respectively. **(c–e)** Luminescence signal from Gluc-RMA, Cluc-RMA, or Nluc-RMA-containing medium. **(c)** CTZ, **(d)** vargulin, and **(e)** furimazine were respectively applied to each RMA medium for evaluating functional orthogonality of detection for each RMA. CTZ specifically reacted with Gluc-RMA in comparison to Cluc-RMA ($P=0.0011$) and Nluc-RMA ($P=0.0015$). Vargulin specifically reacted with Cluc-RMA in comparison to Gluc-RMA ($P<0.0001$) and Nluc-RMA($P<0.0001$). Furimazine specifically reacted with Nluc-RMA in comparison to Gluc-RMA ($P<0.0001$) and Cluc-RMA ($P<0.0001$). One-way ANOVA ($F_{2,9}=17.08$, $P=0.0009$ for CTZ (c), $F_{2,9}=79.57$, $P<0.0001$ for vargulin (d), and $F_{2,9}=44.71$, $P<0.0001$ for furimazine (e)) with Sidak test was used for analysis. **(g)** Nluc-RMA secretion in response to the tdTomato expression. RLU values measured from the culture media reveal the target mRNA-dependent secretion of Nluc-RMA. n=4 independent cultures were analyzed. In comparison of each group with and without the transfection of CAG-tdT, The signal fold change increased over time (5.74 ± 1.47-fold, $P=0.0014$ at 12 h, 29.1 ± 4.74-fold, $P<0.0001$ at 24 h, 68.4 ± 7.46-fold, $P<0.0001$ at 48 h, and 111.4 ± 6.02-fold, $P<0.0001$ at 72 h over the baseline, using two-tailed unpaired t-test using *Benjamini–Hochberg* for FDR adjustment (5%)). *$P<0.05$, **$P<0.01$, ***$P<0.001$, and ****$P<0.0001$. Data is shown as mean ± s.d.

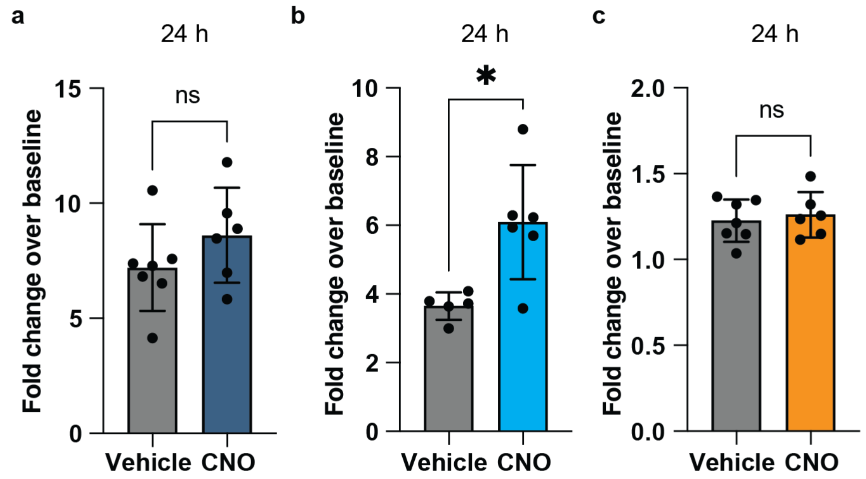


**Extended Data Fig. 8. Multiplexed monitoring of *Arc* expression at 24 h time point by using orthogonal RMA reporters.**

**(a–c)** Relative fold change of plasma bioluminescence over 0 h baseline of **(a)** Gluc-RMA, **(b)** Nluc-RMA, and **(c)** Cluc-RMA at 24 h post-CNO or vehicle administration (P=0.2270 for Gluc, P=0.0111 for Nluc, and P=0.6374 for Cluc-RMAs using two-tailed unpaired t-test). For Gluc- and Cluc-RMAs n=7 (vehicle) and n=6 (CNO) mice were analyzed, while Nluc-RMA used n=5 (vehicle) and n=6 (CNO) mice were analyzed. *P<0.05, ns P>0.05. Data is shown as mean ± s.d.
