## Supplementary data for "Monitoring in vivo transcription with synthetic serum markers"

### Supplementary information

### Table of Contents

**Fig. S1.** Design and screen of sesRNA targeting *TH* mRNAs in vitro.

**Fig. S2.** Design and screen of sesRNA targeting *Fos* mRNAs in vitro.

**Fig. S3.** Design and screen of sesRNA targeting *Arc* mRNAs in vitro.

**Fig. S4.** Histology of brain tissues expressing Fos-targeting sensor.

**Fig. S5.** Immediate response of Arc-targeting sensor by DREADD-induced neuronal activation.

**Fig. S6.** Histology of brain tissues expressing Arc-targeting sensor in the striatum.

**Fig. S7.** Histology of hippocampus tissue slices from the brain expressing three different RMAs.

**Fig. S8.** Histology of striatum tissue slices from the brain expressing three different RMAs.

**Fig. S9.** Histology of hippocampus tissue slices from the brain expressing three different RMAs.

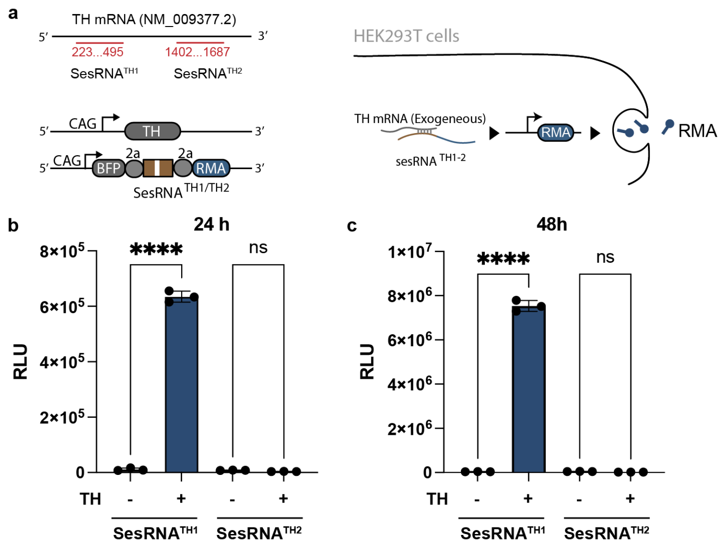


**Supplementary Figure 1. Design and screen of sesRNA targeting *TH* mRNAs in vitro.**
**(a)** The location of the complementary region of the sesRNA within the *TH* mRNA, and the genetic circuit used to test the TH-targeting sensor. The sequences complementary to *TH* mRNA were cloned downstream to the BFP and T2a coding region driven by a CAG promoter. White line in the SesRNA region denotes a single stop codon. **(b, c)** RMA secretion in response to exogenous TH expression. RLU values measured from the culture media showed target mRNA dependent secretion of RMA. n=3 independent cultures were analyzed. For two different TH targeting sensor, in comparison of each group with and without the transfection of CAG-TH from the post-transfection timepoint at 24 h (P<0.00001 for sesRNA (TH1), n.s. P>0.05 for sesRNA (TH2)) and 48 h (P<0.00001 for sesRNA (TH1) and n.s. P>0.05 for sesRNA (TH2)). Data comparisons used unpaired two-tailed multiple t-test with the *Benjamini-Krieger-Yekutieli* for false discovery rate (FDR) adjustment. ****P<0.0001. Data are shown as mean ± s.d.

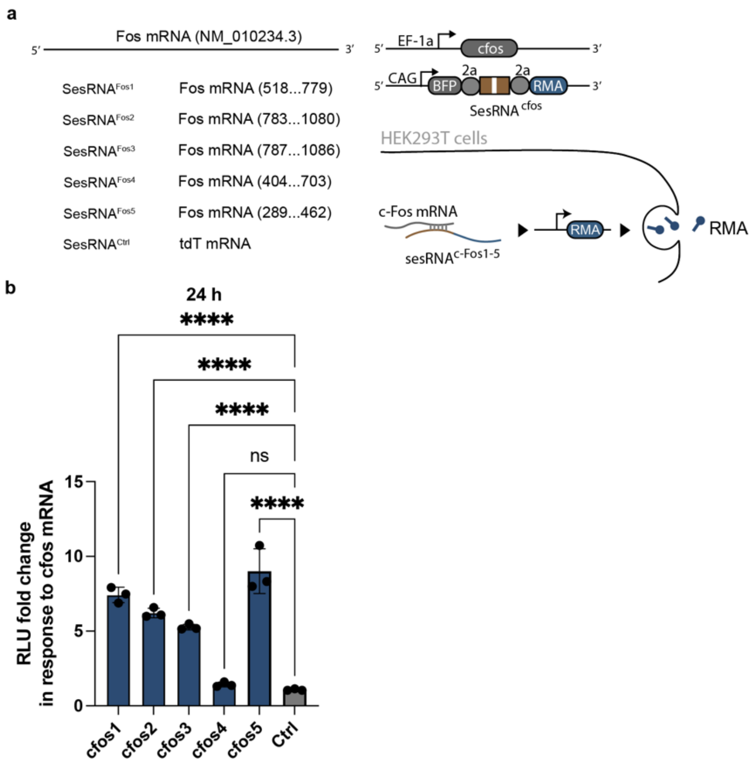


**Supplementary Figure 2. Design and screen of sesRNA targeting *Fos* mRNAs in vitro.** **(a)** The location of the complementary region of the sesRNA within the *Fos* mRNA, and the genetic circuit used to test the Fos-targeting sensor. The sequences complementary to *Fos* mRNA were placed downstream to the BFP and T2a coding region and driven by a CAG promoter. *Fos* mRNA was overexpressed by transfection of plasmid encoding *Fos* downstream of EF-1a promoter (Addgene #193066). White line in the SesRNA region denotes a single stop codon. **(b, c)** Fold activation of *Fos*-targeting RNA sensor in response to exogenous Fos mRNA as compared to the control sensor targeting tdTomato. Data recorded at 24 h after transfection. n=3 independent cultures were analyzed. *tdTomato* mRNA targeting sensor was used in the control (Ctrl) group. ****$p<0.001$ in comparison between *Fos* targeting RNA sensor (Fos1, Fos2, Fos3, and Fos5) and Ctrl sensor, and n.s. ($P>0.05$) between RNA sensor (Fos4) and Ctrl, using one-way ANOVA ($F_{5,12}=69.59$, $P<0.0001$) with Tukey test. ****$P<0.0001$. All data are shown as mean ± s.d.

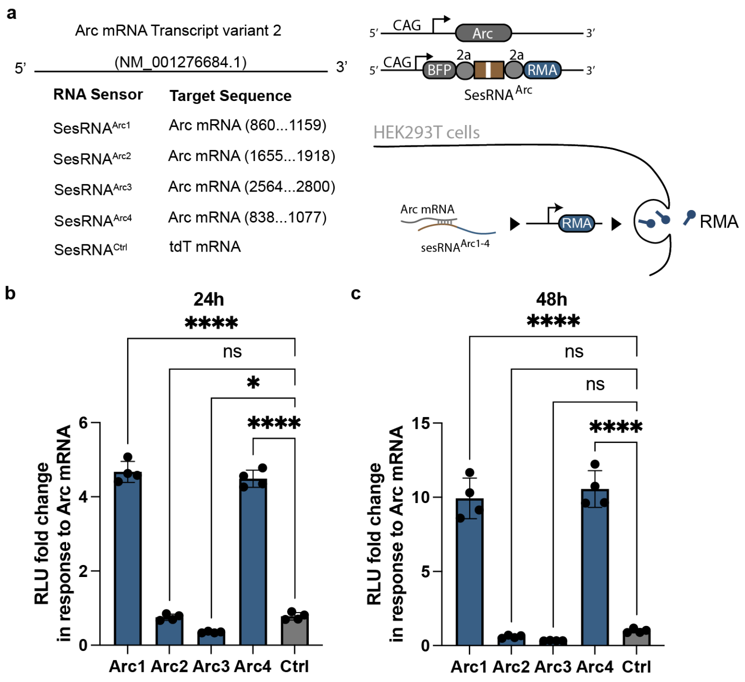


**Supplementary Figure 3. Design and screen of sesRNA targeting *Arc* mRNAs in vitro.** **(a)** The location of the complementary region of the sesRNA within the *Arc* mRNAå and the genetic circuit used to test the Arc-targeting sensor. The sequences complementary to *Arc* mRNA were placed downstream to the BFP and T2a coding region and driven by a CAG promoter. *Arc* mRNA was overexpressed by transfection of plasmid encoding Arc coding sequence downstream of CAG promoter. Arc coding sequence was cloned from mouse cDNA library. White line in the SesRNA region denotes a single stop codon. **(b)** Fold activation of *Arc*-targeting RNA sensor in response to exogenous Fos mRNA as compared to the control (Ctrl) sensor targeting tdTomato. Data recorded at 24 h after transfection. n=3 independent cultures were analyzed. p<0.0001 in comparison between Arc targeting RNA sensor (Arc1 and Arc4) and Ctrl sensor, p=0.0113 in comparison between Arc targeting RNA sensor (Arc3) and Ctrl sensor and n.s. (P>0.05) between RNA sensor (Arc2) and Ctrl, using one-way ANOVA ($F_{4,15}$=626.8, P<0.0001) with Dunnett's test**.** **(c)** Fold activation of Arc-targeting RNA sensor in response to exogenous Fos mRNA as compared to the control (Ctrl) sensor targeting tdTomato at 48 h. n=3 independent cultures were analyzed. tdTomato mRNA targeting sensor was used in the Ctrl group. p<0.0001 in comparison between Arc targeting RNA sensor (Arc1, and Arc4) and Ctrl sensor, and n.s. (P>0.05) between RNA sensor (Arc2 and Arc3) and Ctrl, using one-way ANOVA ($F_{4,15}$=162.4, P<0.0001) with Dunnett's test. *P<0.05, ****P<0.0001, ns P>0.05. All data are shown as mean ± s.d.

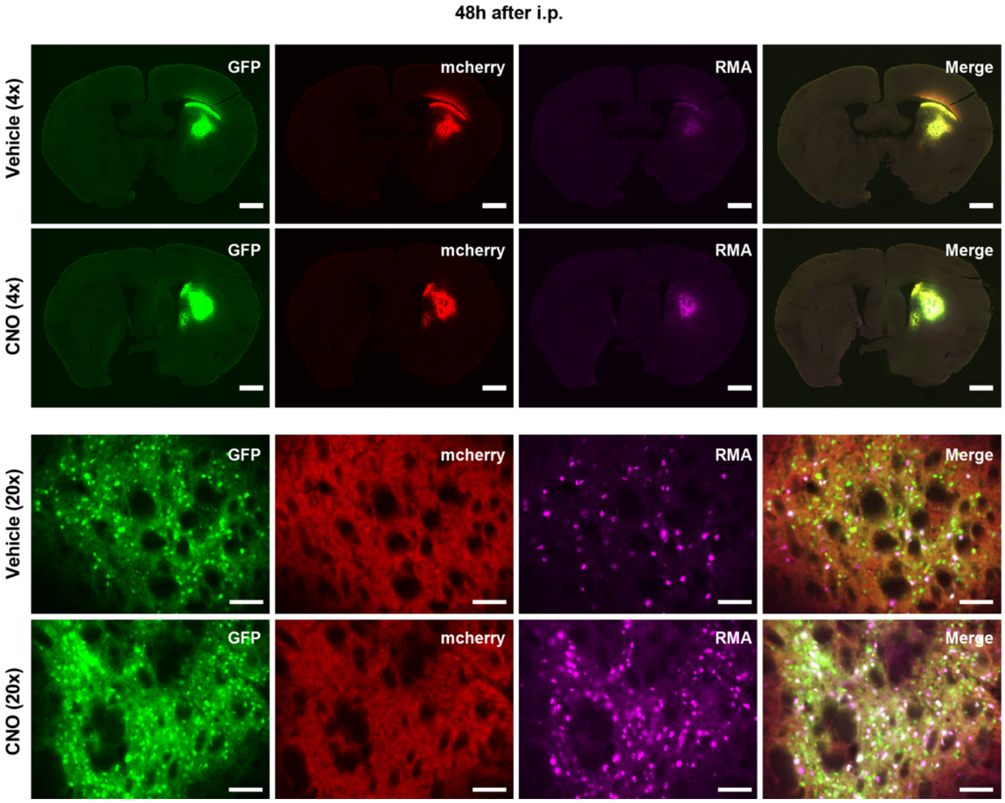


**Supplementary Figure 4. Histology of brain tissues expressing Fos-targeting sensor.** Whole-brain and high magnification images of gene expression of RNA sensor (GFP), DREADD (hM3D(Gq)) receptor (mCherry), and Gluc-RMA (Magenta) in the brain of mice at 48 h after CNO or vehicle (-CNO) injection. Scale bars 1000 μm and 100 μm of the top left (whole brain) and the others (enlarged) images, respectively.

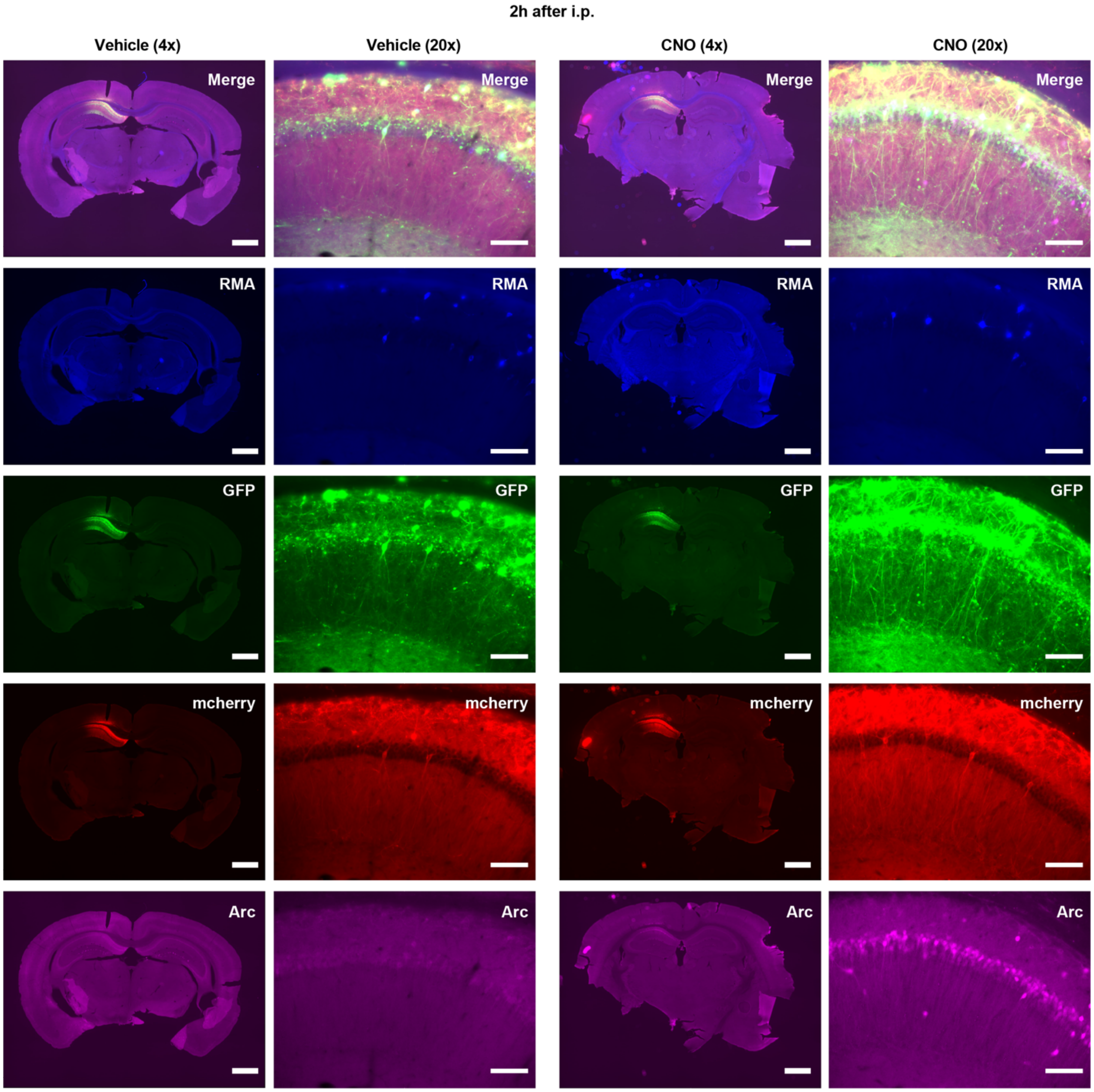


**Supplementary Figure 5. Histology showing Arc induction sensor by DREADD-induced neuronal activation.**

Whole-brain and high magnification images of gene expression of Gluc-RMA (Blue), RNA sensor (GFP), DREADD (hm3D(Gq)) (mCherry), and Arc (Magenta) in the brain 2 h after CNO or vehicle (CNO-) injection. Scale bars 1000 μm and 100 μm of the top left (whole brain) and the others (enlarged) images, respectively.

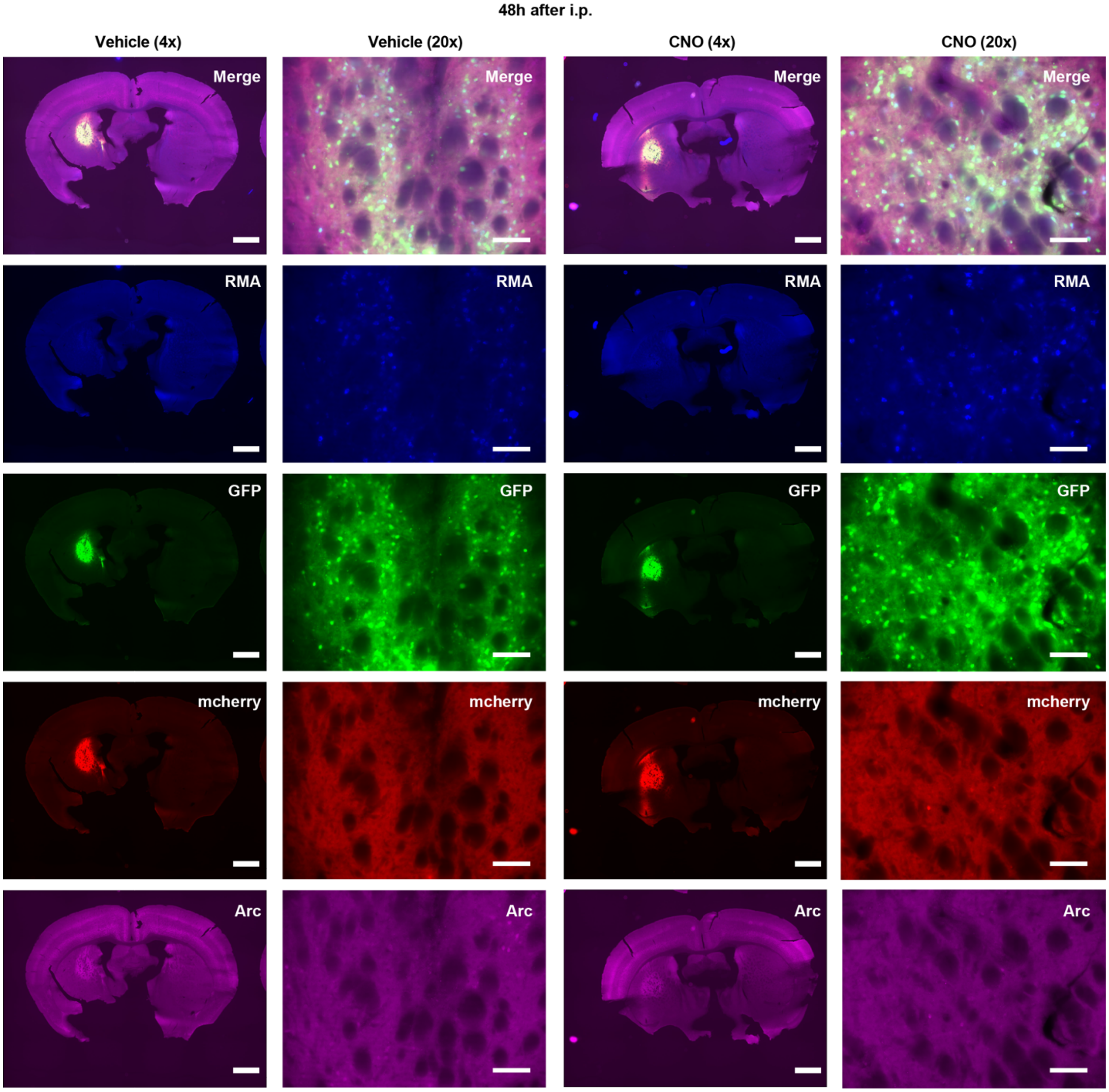


**Supplementary Figure 6. Histology of brain tissues expressing Arc-targeting sensor in the striatum.**

Whole-brain and high magnification images of gene expression of Gluc-RMA (Blue), RNA sensor (GFP), DREADD Gq (mecherry), and Arc (Magenta) in the brain slice at 48 h after intraperitoneal injection (i.p.) from the vehicle (CNO-) mice and CNO-injected mice. AAVs were injected into the lateral side of the sitriatum. Scale bars 1000 μm and 100 μm of the top left (whole brain) and the others (enlarged) images, respectively.

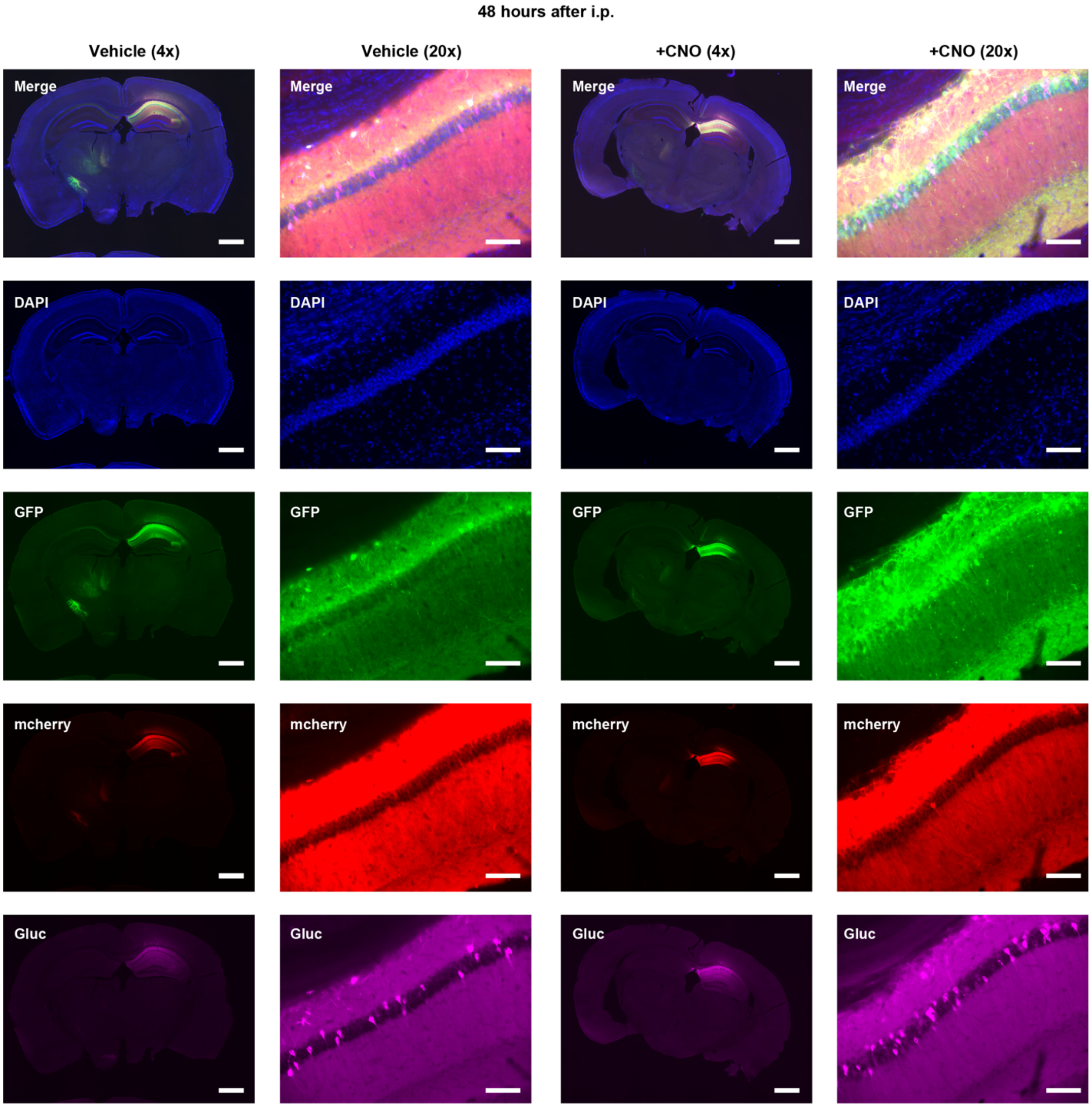


**Supplementary Figure 7. Histology of hippocampus tissue slices from the brain expressing three different RMAs.**
Whole-brain and high magnification images of gene expression of RNA sensor (GFP), DREADD (hM3D(Gq)) (mCherry), and Gluc-RMA (Magenta) in the brain slice at 48 h after intraperitoneal injection (i.p.) from the vehicle mice and CNO-injected mice. AAVs were injected into the lateral side of the hippocampus CA1. Scale bars 1000 μm and 100 μm of the top left (whole brain) and the others (enlarged) images, respectively.

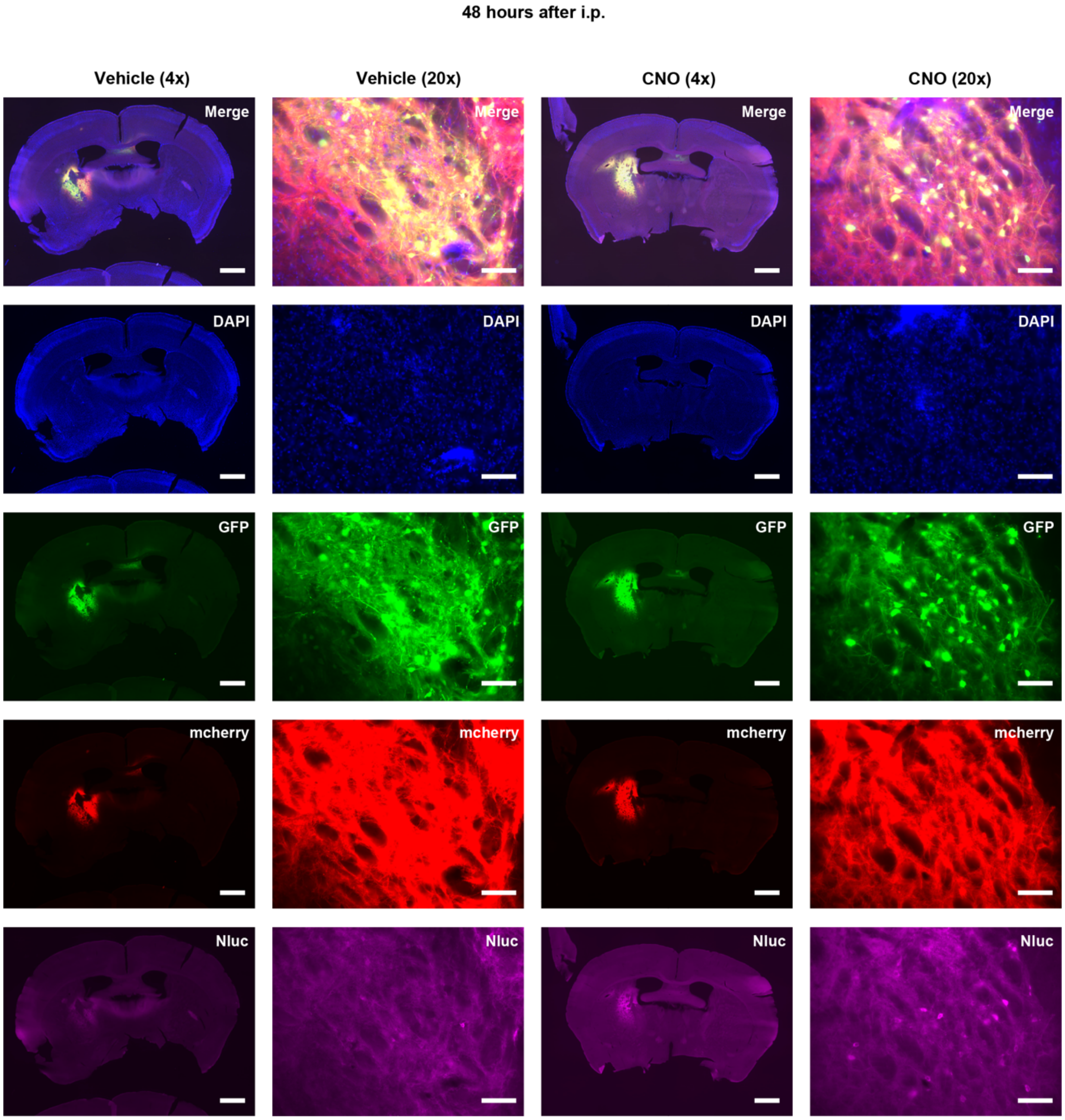


**Supplementary Figure 8. Histology of striatum tissue slices from the brain expressing three different RMAs.**

Whole-brain and high magnification images of gene expression of RNA sensor (GFP), DREADD Gq (mCherry), and Nluc-RMA (Magenta) in the brain slice at 48 h after intraperitoneal injection (i.p.) from the vehicle (CNO-) and CNO-injected mice. AAVs were injected into the lateral side of the striatum. Scale bars 1000 μm and 100 μm of the top left (whole brain) and the others (enlarged) images, respectively.

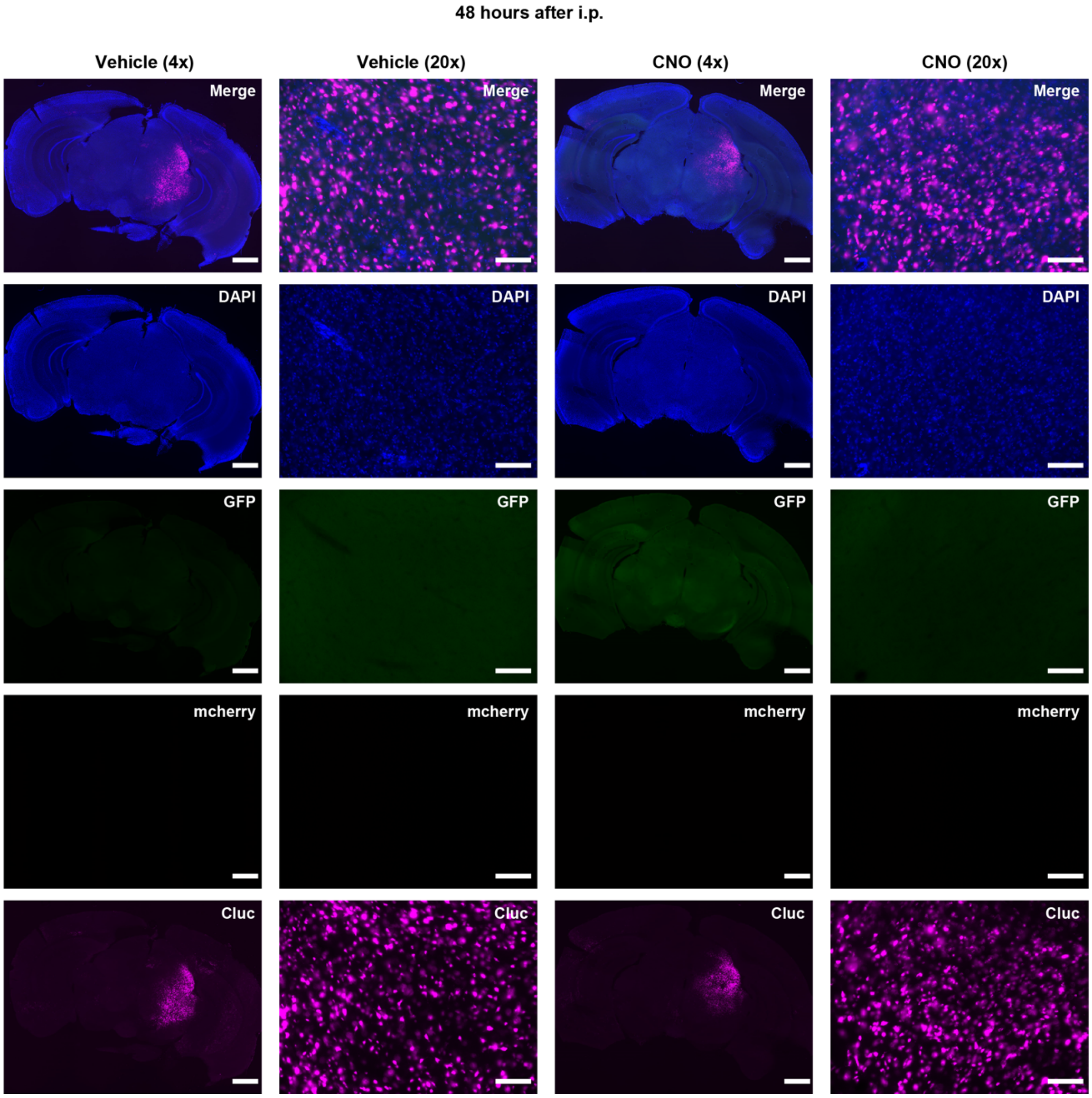


**Supplementary Figure 9. Expression of Cluc in the midbrain.** Whole-brain and high magnification images of gene expression of RNA sensor (GFP), DREADD Gq (mecherry), and Cluc-RMA (Magenta) in the midbrain at 48 h after intraperitoneal injection (i.p.) from the vehicle mice and CNO-injected mice. AAVs were injected into the lateral side of the substantial nigra of the midbrain. Cluc-RMA was stained by using in situ hybridization. Scale bars 1000 μm and 100 μm of the top left (whole brain) and the others (enlarged) images, respectively.

| sesRNA No. | The name of sesRNA | Target mRNA | The number of stop codon (TAG) | Note on construct | Sequence |
|---|---|---|---|---|---|
| 1 | tdT, 1TAG | tdTomato | 1 | Plasmid#192063 was purchased from Addgene | actccaccaggtagtggccgccgtccttcagcttcagggcctggtggatctcgcccttcagcacgccgtc<br>gcgggggtacaggcgctcggtggaggcctcccagcccatagtcttcttctgcattacggggccgtcggggg<br>ggaagttggtgccgcgcatcttcaccttgtagatcagcgtgccgtc |
| 2 | tdT, 2TAG | tdTomato | 2 | | actccaccaggtagtggccgccgtccttcagcttcagggcctggtggatctcgcccttcagcacgccgtc<br>gcgggggtacaggcgctcggtggaggcctcccagcccatagtcttcttctgcattacggggccgtcggggg<br>ggaagttagtgccgcgcatcttcaccttgtagatcagcgtgccgtc |
| 3 | TH1 | Mouse tyrosine hydoroxylase | 1 | | tcagacacccgacgcacagaactgaggagggcagccaggtcgccactgggcacctcgaagcgcacaa<br>agtactccaggtgggggcttcctgccagtggcctctgggcaggccgggtctctaagtagtggattttggcttc<br>aaatgtctcaaacactttcaaagcccgagacagtgaggagggttttgtacccctcaaggagaagagcag<br>gttgagaacagcatttccatccctctcctcgaataccacagcctccaatgggttcccaggttcc |
| 4 | TH2 | Mouse tyrosine hydoroxylase | 1 | | gatgtggttgagaagcagtggaggagggggaccaagagcagcccatcaaagggtccagccacacacatg<br>ggaaagcctctgggccaggaaaggttggagaaggggctgggaactttgggaggcccctggcacctgtaggt<br>ggtaccctatgcatttagctactggcactcagtgcttgggtcagggtgtgcagctcatcctggaccccctct<br>aaggagcgccggatggtgtgaggactgtccagtacatcaatggccagggtgtacgggtcaaacttcacag |
| 5 | Vgat | Mouse vesicular GABA transporter | 1 | Referred to the sequence from Qian et al., 2022 | ggccacatcgaagaagacctggtgccacagcagcttgcgccagagaagacgcaagtggaagaggctgg<br>gcagcaggaagcagaggccggctcccgtgaggctgcccgtgaggcccatgagcagcgc<br>gaagtgtagcacgtagatggccatgagcagcgtgaagaccaccagcgcgcagcgcagcgtcagccccc<br>aggacttaaggcgaccgtcgccaccgtagcaggcggggaagaaggcacgactgccttcct |
| 6 | fos1 | Mouse c-fos | 1 | | aggtcatcggggatcttgcaggcaggtcggtgggctgccaaaataaactccagtttttccttctctttcagc<br>agattggcaatctcagtctgcaacgcagacttctcatcttcaagttcatctgtctccgcttagagtgtatct<br>gtcagctccctcctccgattccggcacttggctgcagccatcttattccgttcccttcggattctccgtttc<br>tcttcctcttcaggagatagctgctctactttgccccttctgccga |
| 7 | fos2 | Mouse c-fos | 1 | | ggaattgctgtgcagaggctcccagtctgctgcatagaaggaaccggacaggtccacatctggcacagag<br>cgggaggtctctcagccactgggcctagatcatgccggaaacaagaagtcatcaaagggttctgccttca<br>gctccacgttgctgatgctcttgactagctccaaggatggcttgggctcagggtcgttgagaaggggcaggg<br>tgaaggcctcctcagactctggggtggaagcctcaggcagacctccagtcaaatccagggaggccacag |

| | | | | |
|---|---|---|---|---|
| 8 | fos3 | Mouse c-fos | 1 | ccccaaggaattgctgtgcagaggctcccagtctgctgcatagaaggaaccggacaggtccacatctgg<br>cacagagcgggaggtctctcagccactgggcctagatcatgccggaaacaagaagtcatcaaagggttc<br>tgccttcagctccacgttgctgatgctcttgactggctccaaggatagcttgggctcagggtcgttgagaag<br>gggcagggtgaaggcctcctcagactctggggtggaagcctcaggcagacctccagtcaaatccaggga |
| 9 | fos4 | Mouse c-fos | 1 | tggcaatctcagtctgcaacgcagacttctcatcttcaagttgatctgtctccgcttggagtgtatctgtcag<br>ctccctcctccgattccggcacttggcagcagccatcttattccgttcccttcggattctccgtttctcttc<br>ctcttcaggagatagctgctctactttgccccttctgccgatgctctgcgctctgcctcctgacacggtctt<br>caccattcccgctctagcgtaagccccagcagactgggtggggagtccgtaaggatggggcgctctggtct |
| 10 | fos5 | Mouse c-fos | 1 | cccagcagactgggtggggagtccgtaaggatggggcgctctggtctgcgatggggccacggaggagacc<br>agagtgggctgcaccagccacagcaggtctaggctggtggagatggctgtcaccgtggggataaagttggc<br>actagagacggacagatctgcgcaaaagtcctgg |
| 11 | Arc1 | Mouse activity-regulated cytoskeleton-associated protein | 1 | tctcctcctcctcagcgtccacatacagtgtctggtacaggtcccgcttgcgccagaggaactggtcgagt<br>ggttcaccctgcttctgcggcagctccagctcccgctgaatagcttcacgggagagtgtaccctcactgta<br>ttgcagaaactccttcttcaactccacccagttcttcaccgagccctgcttcaactcccaccacttcttg<br>gctggcccattcatgtggttctggatctgggacagccaatattcttcagagccacccacctgccgcaggta |
| 12 | Arc2 | Mouse activity-regulated cytoskeleton-associated protein | 1 | cccaagactggccggttgccctgggcggcagctagggtaccagctggatctgctgtgtcagcaggagtggg<br>ctgcatccccggaagttcaggttccctcagcatctctgctttagcatctgccctaggatgtcccctgggttt<br>tggtgcctactttttgttgcctttcagacacttggtttcttcattacccaaagagccctagacactggagca<br>gaatgaggaagccagatcgtgttctgtgttggtctcagagtgtagagggc |
| 13 | Arc3 | Mouse activity-regulated cytoskeleton-associated protein | 1 | gatctgtcagaacatcagtggaattggtctcagaacaccaatagaccaggcagatggaggaaccgcaac<br>aaggcctactcagacagcagacacaagcagctaccagcacaaggtaattggagctagggcttggctgag<br>gtttcagctgggcaatcaccaggcaggtcagggaggttcaaggaacagcctaggagccaaggcctggggt<br>ctcctgggactgggcttgacctggcaggcc |
| 14 | Arc4 | Mouse activity-regulated cytoskeleton-associated protein | 1 | cttctgcggcagctccagctcccgctgaatggcttcacgggagagtgtaccctcactgtattgcagaaact<br>ccttcttgaactccacccagttcttcaccgagccctgcttgaactcccaccacttcttggctagcccatt<br>catgtggttctggatctgggacagccaatattcttcagagccacccacctgccgcaggtactcttccaggt<br>ggctcaggaactcccgtgggtcctcgaac |

| Plasmids | Figure index | Construct (backbone) | Note on construct |
|---|---|---|---|
| 1 | Fig. 2 | CAG-tdTomato | Addgene#83029 |
| 2 | Fig. 2a, b, c | CAG-BFP-P2A-sesRNA(tdT)-T2A-Gluc-RMA | Backbone from Addgene#192063. |
| 3 | Supplementary Fig. 1 | CAG-BFP-P2A-sesRNA(TH1/TH2)-T2A-Gluc-RMA | sesRNA was cloned into the plasmid No.2 |
| 4 | Supplementary Fig. 1 | CAG-TH | Th insert was coned from addgene#105988). Backbone from the Plasmid 2 |
| 5 | Supplementary Fig. 2 | CAG-BFP-P2A-sesRNA(fos1/fos2/fos3/fos4/fos5)-T2A-Gluc-RMA | sesRNA was cloned into the plasmid No.2 |
| 6 | Supplementary Fig. 2 | pHAGE-Fos-mcherry | Addgene#193066 |
| 7 | Supplementary Fig. 3 | CAG-BFP-P2A-sesRNA(Arc1/Arc2/Arc3/Arc4)-T2A-Gluc-RMA | sesRNA was cloned into the plasmid No.2 |
| 8 | Supplementary Fig. 3 | CAG-Arc | Th insert was coned from N2a cell-derived cDNA library. Backbone from the Plasmid 2 |
| 9 | Extended data Fig. 1 | CMV-ADAR1p150 | Addgene#117927 |
| 10 | Extended data Fig. 1 | CAG-ADAR2-sesRNA(tdT)-Gluc-RMA-W3SL | Backbone from Addgene#102069, GFP was replaced with Gluc-RMA. |
| 11 | Fig. 2d, e, f | CAG-BFP-P2A-sesRNA(tdT)-T2A-tTA2 | Addgene#192070 |
| 12 | Fig. 2d, e, f | CAG-BFP-P2A-sesRNA(tdT_2TAG)-T2A-tTA2 | Point mutation was inserted into sesRNA(tdT) in plasmid#5 |
| 13 | Fig. 2d, e, f, Fig. 3, 4, 5, 6, 7 | pAAV-TRE3g-Gluc-RMA | Backbone from Addgene#192064, mNeonGreen was replaced with Gluc-RMA. |
| 14 | Extended data Fig. 2 | CAG-Gluc-RMA | Backbone from Addgene#83029, tdTomato was replaced with Gluc-RMA. |
| 15 | Fig. 3 | pAAV-hSyn1-GFP-P2A-sesRNA(tdT)-T2A-tTA2 | Backbone from Addgene #189629 |
| 16 | Fig. 3 | pAAV-CAG-tdT | Backbone from Addgene #189629 |
| 17 | Fig. 3, 4, 5, 6, 7 | pAAV-hSyn1-Cluc-RMA | Addgene #189624 |
| 18 | Fig. 4 | pAAV-hSyn1-GFP-P2A-sesRNA(TH)-T2A-tTA2 | Backbone from Addgene #189629 |
| 19 | Extended data Fig. 4 | pAAV-hSyn1-GFP-P2A-sesRNA(vgat)-T2A-tTA2 | Backbone from Addgene #189629 |
| 20 | Fig. 5 | pAAV-hSyn1-GFP-P2A-sesRNA(cfos)-T2A-tTA2 | Backbone from Addgene #189629 |
| 21 | Extended data Fig. 5, Fig. 6 | pAAV-hSyn1-GFP-P2A-sesRNA(Arc)-T2A-tTA2 | Backbone from Addgene #189629 |
| 22 | Fig. 6 | pAAV-TRE3g-Nluc-RMA | Backbone plasmid No. 10. Gluc was replaced with Nluc. |
